# An Arf6- and caveolae-dependent pathway links hemidesmosome remodeling and mechanoresponse

**DOI:** 10.1101/126151

**Authors:** Naël Osmani, Julien Pontabry, Jordi Comelles, Nina Fekonja, Jacky G. Goetz, Daniel Riveline, Elisabeth Georges-Labouesse, Michel Labouesse

## Abstract

Hemidesmosomes (HDs) are epithelial-specific cell-matrix adhesions, which stably anchor the intracellular keratin network to the extracellular matrix. Although their main role is to protect the epithelial sheet from external mechanical strain, how HDs respond to mechanical stress remains poorly understood. Here we identify a pathway essential for HD remodeling, and outline its role with respect to α6β4 integrin recycling. We find that α6β4 integrin chains localize to the plasma membrane/caveolae and Arf6+ endocytic compartments. Based on FRAP and endocytosis assays, integrin recycling between both sites requires the small GTPase Arf6 and its co-regulators GIT1/βPIX, but neither Caveolin1 (Cav1) nor Cavin1. Strikingly, when keratinocytes are stretched or hypo-osmotically shocked, α6β4 integrin accumulates at cell edges, whereas Cav1 disappears from it. This process, which is isotropic relative to the orientation of stretch, depends on Arf6, Cav1 and Cavin1. We propose that mechanically-induced HD growth involves the isotropic flattening of caveolae (known for their mechanical buffering role) associated with integrin diffusion and turnover.

## Introduction

The physical interaction of cells with their environment is essential for their function and the maintenance of tissue architecture and homeostasis. Cell adhesion to the ECM provides a link and allows the cell to probe the physical and chemical properties of its microenvironment (Balaban et al., 2001; Galbraith et al., 2002; Riveline et al., 2001). Two classes of cell-ECM adhesion complexes, with their distinctive integrins and adaptor proteins, enable cell attachment and migration: focal adhesions (FAs) and epithelial-specific hemidesmosomes (HDs) (Hopkinson et al., 2014; Kim et al., 2011). While the polarized maturation of FAs under mechanical cues by further recruitment of adaptor proteins is well described (Moore et al., 2010; Parsons et al., 2010), our knowledge of the mechanisms controlling HD remodeling remains incomplete. In particular, the input of mechanical forces has not been investigated in vertebrates.

The HD-specific integrin α6β4 links components of the extracellular basement membrane, in particular the epithelial-specific laminin-332, to the intracellular plectin and BPAG1e, which in turn anchor the keratin cytoskeleton (Litjens et al., 2005; Walko et al., 2011; Rezniczek et al., 1998). HD-mediated adhesion to the extracellular basement membrane, coupled to the high deformability of the keratin cytoskeleton protect the epithelial layer from mechanical strain (Herrmann et al., 2007; Osmani and Labouesse, 2015). Indeed, patients with defective HDs due to inherited (epidermolysis bullosa) or autoimmune skin diseases develop severe blistering syndromes (Iwata and Kitajima, 2013; Sawamura et al., 2010).

Vertebrate HDs undergo dynamic remodeling during keratinocyte migration and wound healing (Osmani and Labouesse, 2015). Biochemical and cellular analyses have highlighted the contribution of β4 integrin phosphorylation in regulating HD disassembly (Frijns et al., 2012; Frijns et al., 2010; Germain et al., 2009; Rabinovitz et al., 2004; Wilhelmsen et al., 2007) [for reviews, see refs (Litjens et al., 2006; Hopkinson et al., 2014; Osmani and Labouesse, 2015; Walko et al., 2014)]. In *C. elegans*, HD-like structures (CeHDs), which attach the epidermis to the cuticle apically and to an ECM shared with the underlying muscles basally, must also be remodeled during embryonic elongation (Zhang and Labouesse, 2010). We recently established that muscle contractions contribute to CeHD remodeling by inducing an epidermal mechanotransduction pathway that stimulates intermediate filament phosphorylation and reorganization (Zhang et al., 2011).

To identify the mechanisms controlling HD remodeling in vertebrates and assess how tension might impact on the process, we implemented a genetic screen inspired from the strategy previously used in *C. elegans* (Zahreddine et al., 2010). We identify several novel regulators of HD turnover, and demonstrate that external mechanical cues drive their remodeling.

## Results

### Identification of new genes involved in HD remodeling

To identify genes controlling HD biogenesis, we relied on the observation that genetic ablation of HD components increases keratinocyte migration speed by two-fold (Osmanagic-Myers et al., 2006; Raymond et al., 2005; Rodius et al., 2007; Seltmann et al., 2013). Specifically, we used stable shRNA expression in the human keratinocytes cell line HaCaT to generate cells with approximately 50% less plectin than control cells. The stable clone shPlec#6 migrated 1.7-fold faster in a scratch assay and had partially defective HDs (Fig. S1A-D). We subsequently examined the migration properties of such cells after silencing human homologs of the genes promoting CeHD biogenesis (Fig. S1F). We expected that knocking-down such genes should further increase the migration speed of shPlec#6 cells to reach that of plectin-KO cells (Fig. S1A,E). Among the 30 genes screened, the Maternal Embryonic Leucine Zipper kinase (MELK) the p21-activated kinase 1 (PAK1) and the Rac/Cdc42 PAK-interacting exchange factor (βPIX) had the strongest effect. Since βPIX and PAK1 form a complex, we decided to focus on those proteins (Frank and Hansen, 2008; Rosenberger and Kutsche, 2006). In addition, although the G protein–coupled receptor kinase–interacting ArfGAP 1 (GIT1) and the small GTPase Rac1 did not clearly affect migration speed, we included them in our study, because βPIX is a Rac Guanine Nucleotide Exchange Factor (Rosenberger and Kutsche, 2006) that interacts with GIT1 and PAK1 (Frank and Hansen, 2008).

### βPIX, PAK1, Rac1 and GIT1 regulate HD formation

To confirm the results from the screen, we carefully analyzed HDs shortly after spreading (1h) and after 72h to observe immature and mature HDs respectively. After 72h, we observed growing HDs on the very edge of cells with well-defined keratin bundles associated to small HD integrin clusters containing plectin (Fig. 1A, arrowheads), and more compact HDs towards the center of the cell (Fig. 1A, above dashed line). We asked whether the HD area at the basal plasma membrane would depend on the spreading time. To do so, we quantified the relative HD area of cells spread for 1h on polylysine-coated circular patterns of 30 μm diameter, or for 1h or 72h on non-patterned surfaces; note that based on the HaCaT migration speed measured for our mini-screen (5∼10 μm/h) cells can spread but not migrate within 1 hour. We observed that HDs occupied a similar area, ranging from 25 to 32% of the basal plasma membrane in all three conditions (Fig. 1B). To determine whether the aforementioned proteins had a direct effect on HD spatial distribution, we depleted them and examined the localization of the α6 integrin subunit (ITGA6) and plectin in wild-type HaCaT cells. We observed that GIT1, βPIX, Rac1 and PAK1 depletion decreased the HD amount at the basal plasma membrane by 75 to 95% after 72h on non-patterned surfaces, and that only GIT1 and βPIX depletion significantly reduced HDs at the basal plasma membrane after 1 hour spreading (Fig. 1C-E’). These observations suggest that GIT1 and βPIX depletion directly affected the localisation of HD integrins at the basal membrane, whereas Rac1 and PAK1 depletion may affect them indirectly or at a later step. Previous work from our group has shown that in *C. elegans* PAK1 acts downstream of Rac1 and is required to reorganize the intermediate filament cytoskeleton in order to promote CeHD remodeling (Zhang et al., 2011). Consistent with this notion, the keratin cytoskeleton appeared disorganized with thin filaments, especially at the cell edge, and was not well organized in a large and dense network as observed in control cells (Fig. 1F-F’). To confirm that keratins had an abnormal organization in PAK1 deficient cells, we used a method adapted from the physics of liquid crystals to quantify their degree of organization through nematic order. We observed a significant disorganization, which, for PAK1-depleted cells was equivalent to ITGA6-depleted cells lacking HDs (Fig. S1G), consistent with recent observation made for keratin-13 in plectin-depleted epidermoid carcinoma cell line (Moch et al., 2016). To test whether the protein-protein interaction domains and the enzymatic activities of the GIT1/βPIX/Rac1/PAK1 signaling module are required for proper HD formation we transfected mutant forms lacking these domains in HaCaT cells. Specifically, we found that a deletion of the GIT1 ArfGAP domain (ΔGAP), which controls the GTPase activity of the trafficking regulator Arf6 (de Curtis, 2001; Donaldson and Jackson, 2011), significantly reduced the HD amount (Fig. S2A,B), that the βPIX domains RacGEF (ΔDH) and SH3 (interacting with PAK1; SH3) are required for HD formation (Fig. S2A,C). Likewise, a dominant-negative Rac1(T17N) affected HD formation, suggesting that the GTPase function of Rac1 is required (Fig. S2A,D). Finally, a kinase-dead PAK1(K299R) as well as a constitutively active PAK1(T423E) forms affected HD formation (Fig. S2A,E), suggesting that PAK1 activity is required and should be tightly regulated.

**Figure 1.**
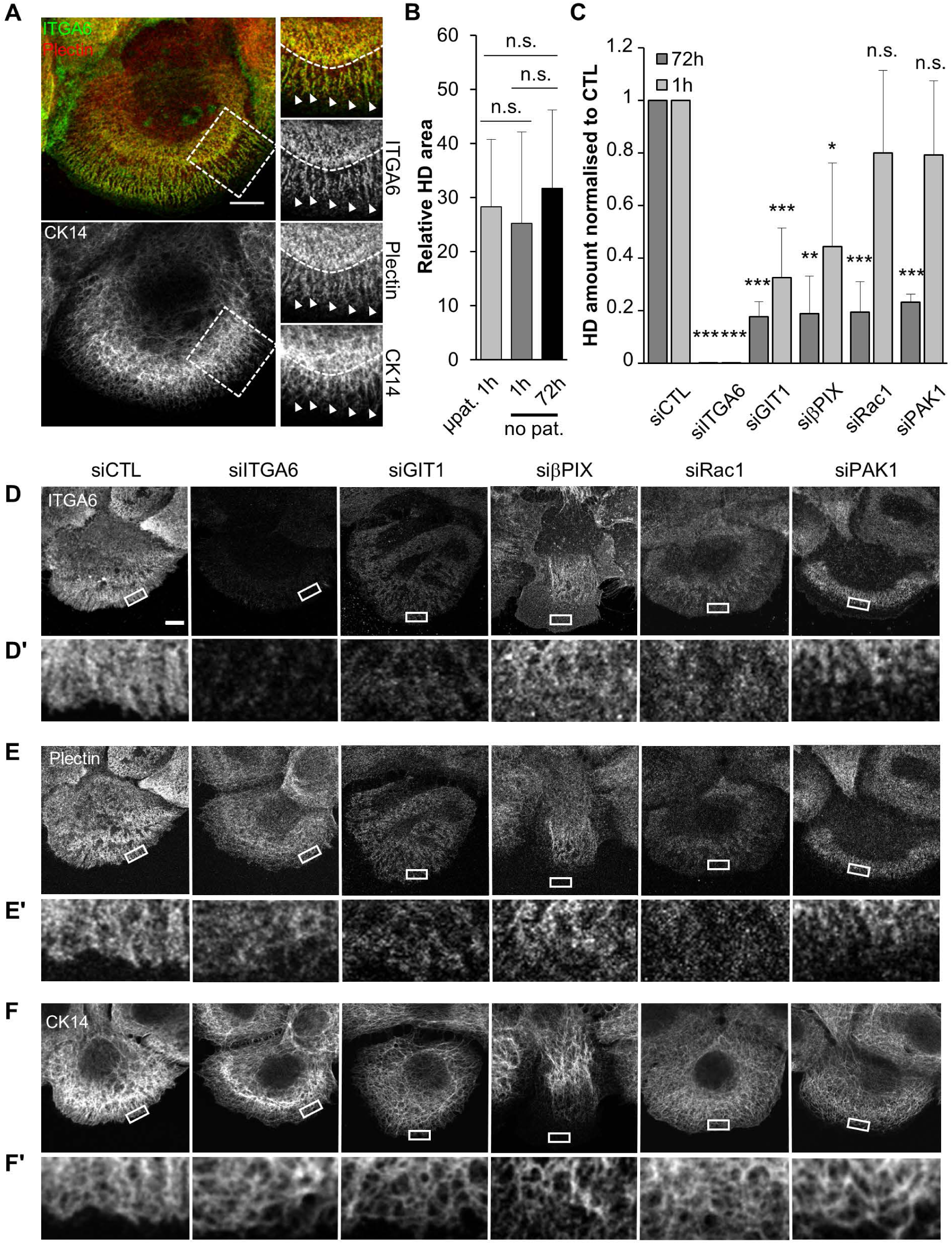
GIT1, βPIX, PAK and Rac1 are required for proper HD organisation. (A) Confocal plane of the basal plasma membrane of HaCaT cells fixed after 72h on non-patterned surfaces and immunostained for ITGA6, Plectin and keratin 14 (CK14). Scale bar=10 μm. (B) Quantification of the HD area at the basal membrane of HaCaT cells spread for 1h on circular patterns of 30 μm in diameter coated with poly-lysine (μpat.) or for 1h or 72h on non-patterned coverslips (no pat.). (C) Quantification of the amount of HDs at the basal membrane of siRNAs transfected cells spread on non-patterned surfaces for 1h or 72h. (D-F’) Confocal plane of the basal plasma membrane of HaCaT cells fixed after 72h on non-patterned surfaces, transfected with control or the indicated siRNAs and immunostained for ITGA6 (A), Plectin (B) or keratin 14 (CK14) (C); brightness and contrast are unmodified. (D’, E’ and F’) Magnifications of the areas boxed in (D, E and R); brightness and contrast have been boosted for clarity. Scale bar=10 μm.

**Figure 2.**
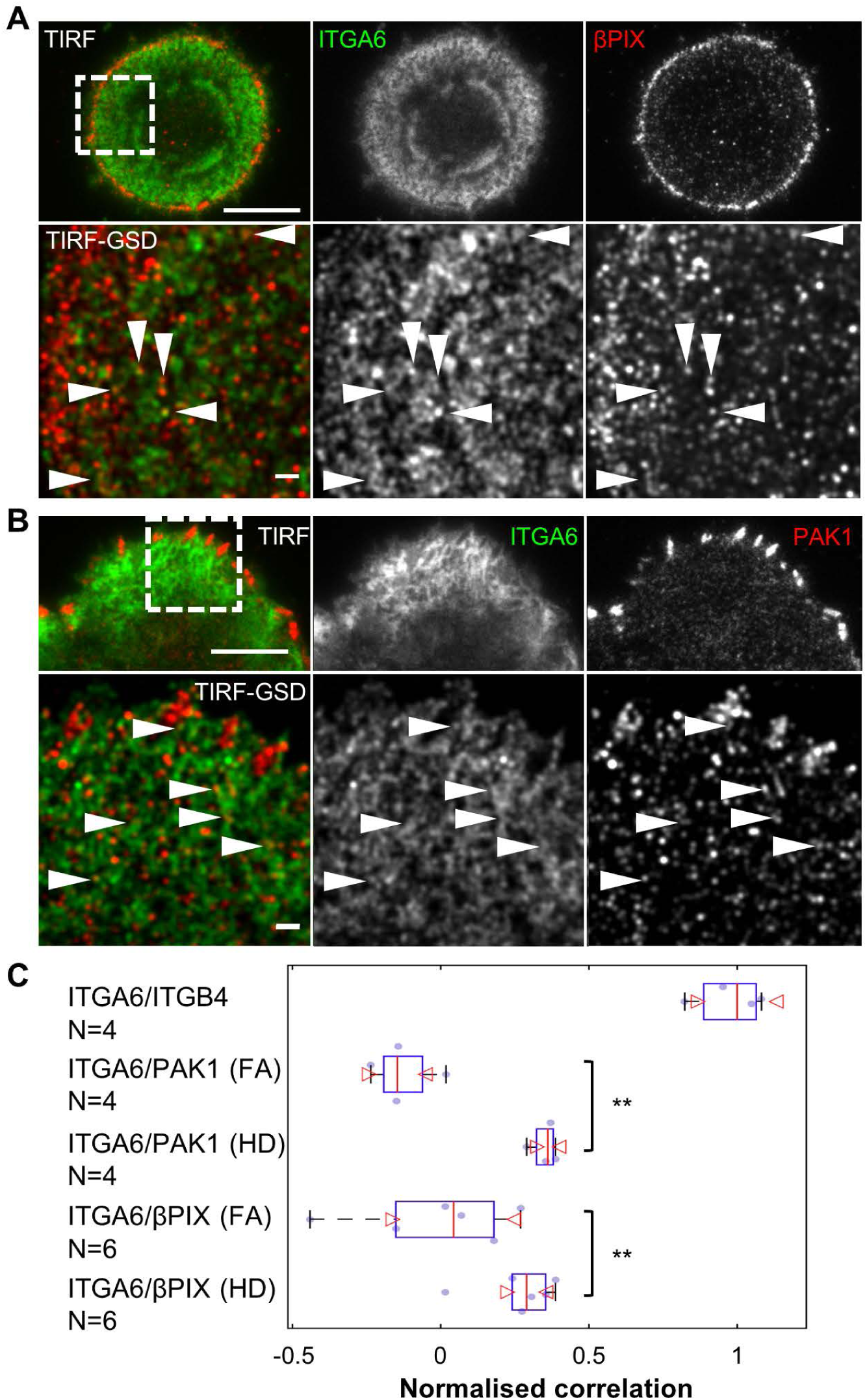
Discrete pools of βPIX and PAK1 colocalise with ITGA6 at the basal membrane. (A-B) HaCaT cell stained for ITGA6 and (B) βPIX or (C) PAK1. Upper panel, TIRF image, scale bar=10 μm; lower panel, GSD-TIRF image (pixel size = 20 nm), scale bar=1 μm. Arrowheads highlight colocalisations. (C) Mean correlation of both channels measured in single GSD images for each pair of markers. Values are normalised to the mean correlation coefficient of ITGA6/ITGB4 GSD images used as positive control (see methods for details).

Although GIT1, βPIX and PAK1 also regulate FA morphology and dynamics in fibroblasts (Deakin and Turner, 2008; Rosenberger and Kutsche, 2006), HD defects did not result from FA defects, since the latter were essentially normal in GIT1/βPIX/PAK1-depleted cells (Fig. S3A-B). Reciprocally, affecting FA formation by inhibiting FAK activity with PF-228 (Schaller, 2010) did not affect HDs (Fig. S3C-D). Likewise, RNAi-mediated knock-down of β1 integrin (ITGB1), one of the major adhesion receptor in keratinocyte FAs (Hopkinson et al., 2014), did not affect HDs (Fig. S3E). Thus, we conclude that GIT1, βPIX, Rac1 and PAK1 directly affect HD remodeling independently of FAs.

To support the functional analysis described above, we examined the subcellular localization of these factors. In addition to their localization at FAs, endogenous βPIX and PAK1 were present at the basal plasma membrane in regions coinciding with HDs marked by ITGA6 (Fig. 2). Indeed, using TIRF illumination and GSD nanoscopy, we found that ITGA6 staining partially correlated with βPIX and PAK1 at the basal plasma membrane (R=0.29 and 0.36 respectively), but was not observed at FAs, which rely on other integrins (Fig. 2).

Collectively, these data show that cells defective for GIT1/βPIX/Rac1/PAK1 have abnormal ITGA6 and keratin distribution, showing that these proteins promote HD formation.

### Arf6, GIT1/βPIX and caveolae control HD biogenesis

The observation that the ArfGAP domain of GIT1 was required for proper HD formation raised the possibility that it acts in part by controlling Arf6 dependent trafficking of α6β4 integrin. In particular, in other cell types, Arf6 regulates FA-mediated adhesion by promoting β1 integrin recycling (Donaldson and Jackson, 2011). Hence, we carefully examined α6β4 integrin localization and found that it was enriched in intracellular compartments (ICs) located 1 μm above HDs (Fig. 3A,A’). These ICs also contained tagged Arf6 and GIT1, endogenous βPIX and PAK1 and tagged Rac1 (Fig. 3B-H). Colabeling Arf6^+^/ITGA6^+^ ICs with trafficking markers suggested that they correspond to recycling endosomes (RE), which is consistent with the known role of Arf6 (Aikawa and Martin, 2003; Powelka et al., 2004). Indeed, less than 20% ICs contained the early endosome marker EEA1, whereas 50% of them had the RE marker Rab11 (Fig. 3B,F). Integrin-positive ICs also contained Cav1 (75%), a marker for caveolae (Ω-shaped membrane invagination of ∼100nm in diameter) (Parton and del Pozo, 2013), previously associated with α6β4 in lipid rafts (Gagnoux-Palacios et al., 2003) (Fig. 3B,G-H). This colocalization was further confirmed by immunogold labeling which showed vesicles containing endogenous ITGA6 and Cav1 close to the plasma membrane (Fig. 3I) or within larger compartments 500 nm below the plasma membrane (Fig. 3J). In addition, we observed endogenous ITGA6 and Cav1 colocalization at the basal plasma membrane by TIRF-GSD nanoscopy and measured a normalized correlation for colocalization significantly higher than for a negative control corresponding to FAs marked by βPIX or PAK1 (Fig. 4A-B). Immuno-gold labeling confirmed that ITGA6 clusters were localized either close to or just at the edge of Cav1+ caveolae (Fig. 4C).

**Figure 3.**
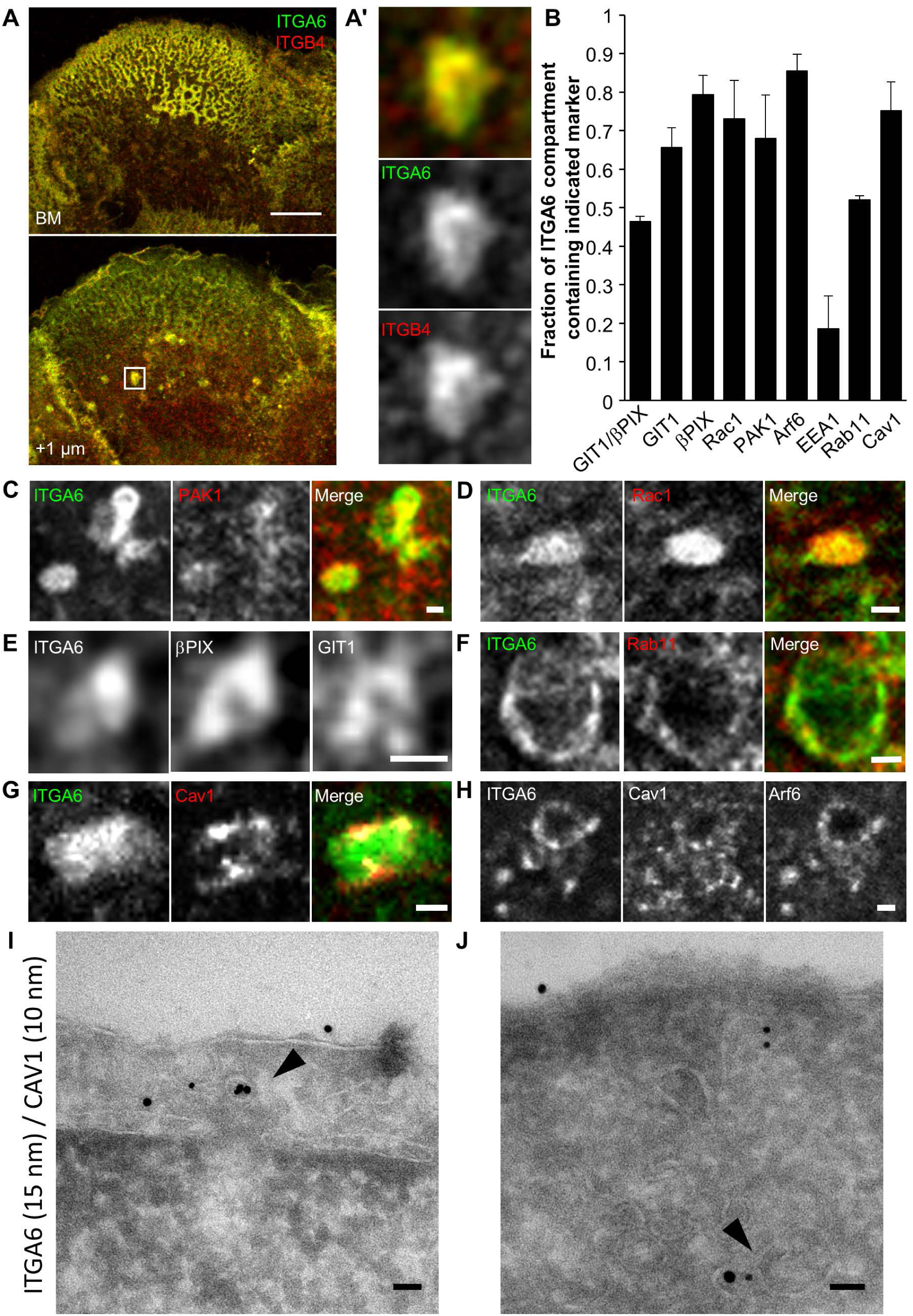
HD integrins are located in Arf6-intracellular compartments containing Cav1, GIT1, βPIX, PAK1 and Rac1. (A) Confocal plane of the basal membrane or +1 μm above basal membrane of HaCaT cells stained for ITGA6 and ITGB4. Scale bar 10 μm. (A’) 10x magnification of the areas boxed in (A). (B) Quantification of the presence of various markers within ITGA6-containing intracellular compartments. N: GIT1/βPIX=21, GIT1=32, βPIX=46, Rac1=35, PAK1=20, Arf6=46, EEA1=16, Rab11=22, Cav1=43 cells. (C-H) Close-up on ICs in a confocal section +1 pm above the basal plasma membrane of cells co-immunostained for ITGA6 and (C) endogenous PAK1; (D) expressing EGFP-Rac1(wt); (E) endogenous βPIX and expressing FLAG-GITI(wt) and; (F) expressing RFP-Cav1(wt); (G) for endogenous Rab11, (H) endogenous Cav1 and expressing EGFP-Arf6. Magnification: (C-D) 3X; (E) 8X; (F) 10X; (G) 5X; (Η) 3X respectively. Scale bar=1 μm, (l-J) Immuno-electron micrographs of HaCaT cells after staining with antibodies against ITGA6 (15 nm gold beads) and Cav1 (10 nm gold beads) showing colocalization of ITGA6 and Cav1 in vesicles close to the plasma membrane (J, arrow) or 500 nm below (I, arrow). Scale bar=50 nm.

**Figure 4.**
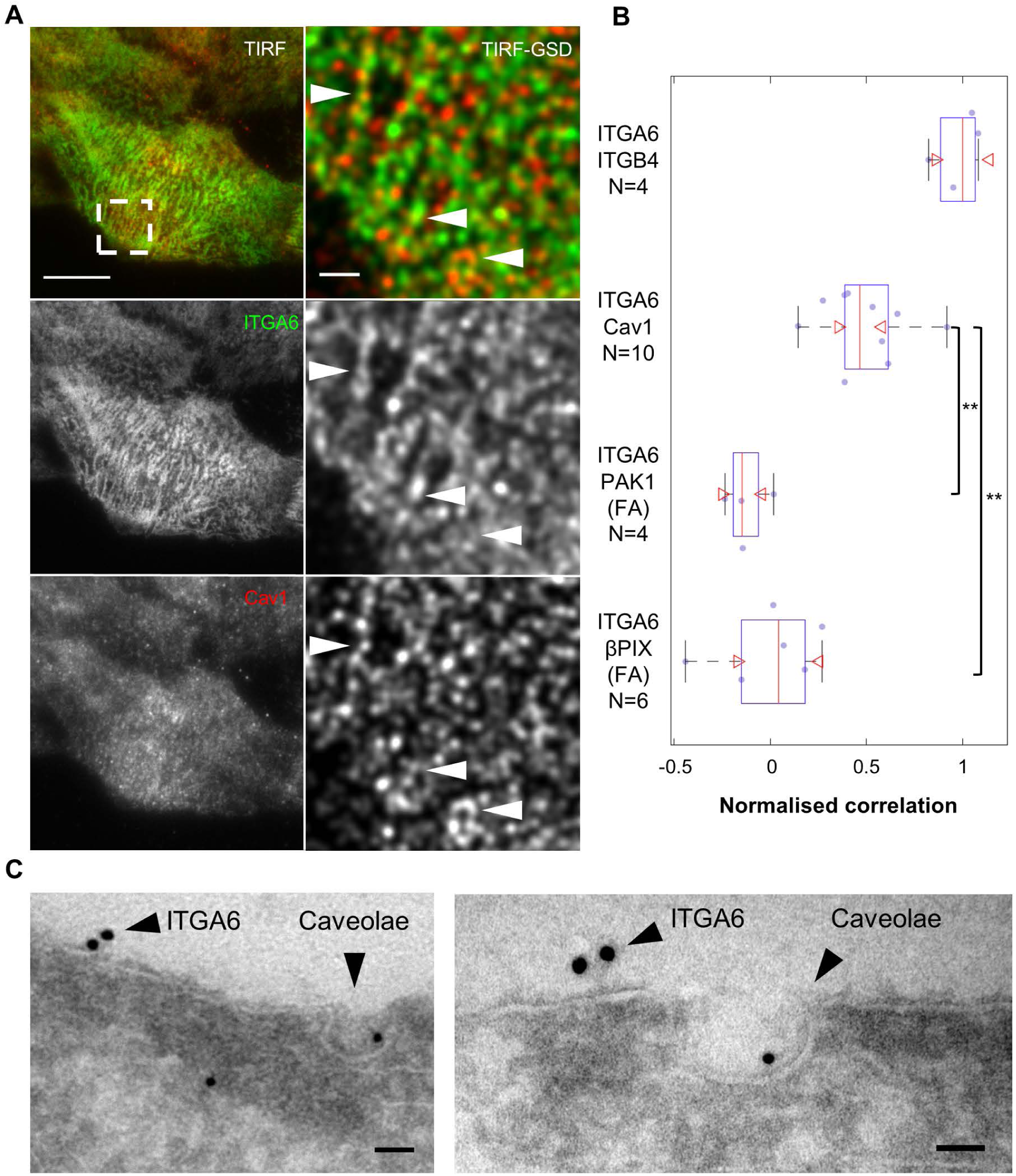
Cav1 colocalises with ITGA6 at the basal membrane. (A) HaCaT cell stained for ITGA6 and Cav1. Left panel: TIRF image, scale bar=10 μm right panel: GSD-TIRF image (pixel size = 20 nm), scale bar=1 μm. Arrowheads highlight colocalisation. (B) Mean correlation of both channels measured in single GSD images for each pair of markers. Values are normalised to the mean correlation coefficient of ITGA6/ITGB4 GSD images used as positive control, and compared to the correlation of ITGA6 at focal adhesions marked by PAK1 or βPIX taken as a negative control (see methods for details). (C) Immuno-electron micrographs of the plasma membrane of HaCaT cells after staining with antibodies against ITGA6 (15 nm gold beads) and Cav1 (10 nm gold beads). Scale bar=50 nm.

To assess whether Arf6- and Cav1 control HD biogenesis, we affected their function using siRNAs or mutant forms. Arf6 depletion induced a strong loss of HDs at the basal plasma membrane, associated with the same thin and less dense IF network as observed when GIT1 or βPIX are depleted (Fig. 5A-B). Likewise, dominant-negative Arf6(T27N)-expressing cells reduced the HD area to the same extent as siRNA-Arf6 (Fig. 5C-D) and fewer ITGA6/Arf6 double positive ICs (Fig. 5E), suggesting that integrin recycling was impaired. The constitutively-active Arf6(Q67L) form induced mild HD defects with ITGA6 clustering centrally with Arf6 at the basal plasma membrane in 57/63 cells (Fig. 5C,F-F’) without any associated plectin or keratins (data not shown) suggesting a misregulation of integrin recycling. Nontransfected cells had the same amount of HDs at the basal plasma membrane as Arf6-expressing cells, suggesting that Arf6 overexpression did not affect HD organization (Fig. 5C).

**Figure 5.**
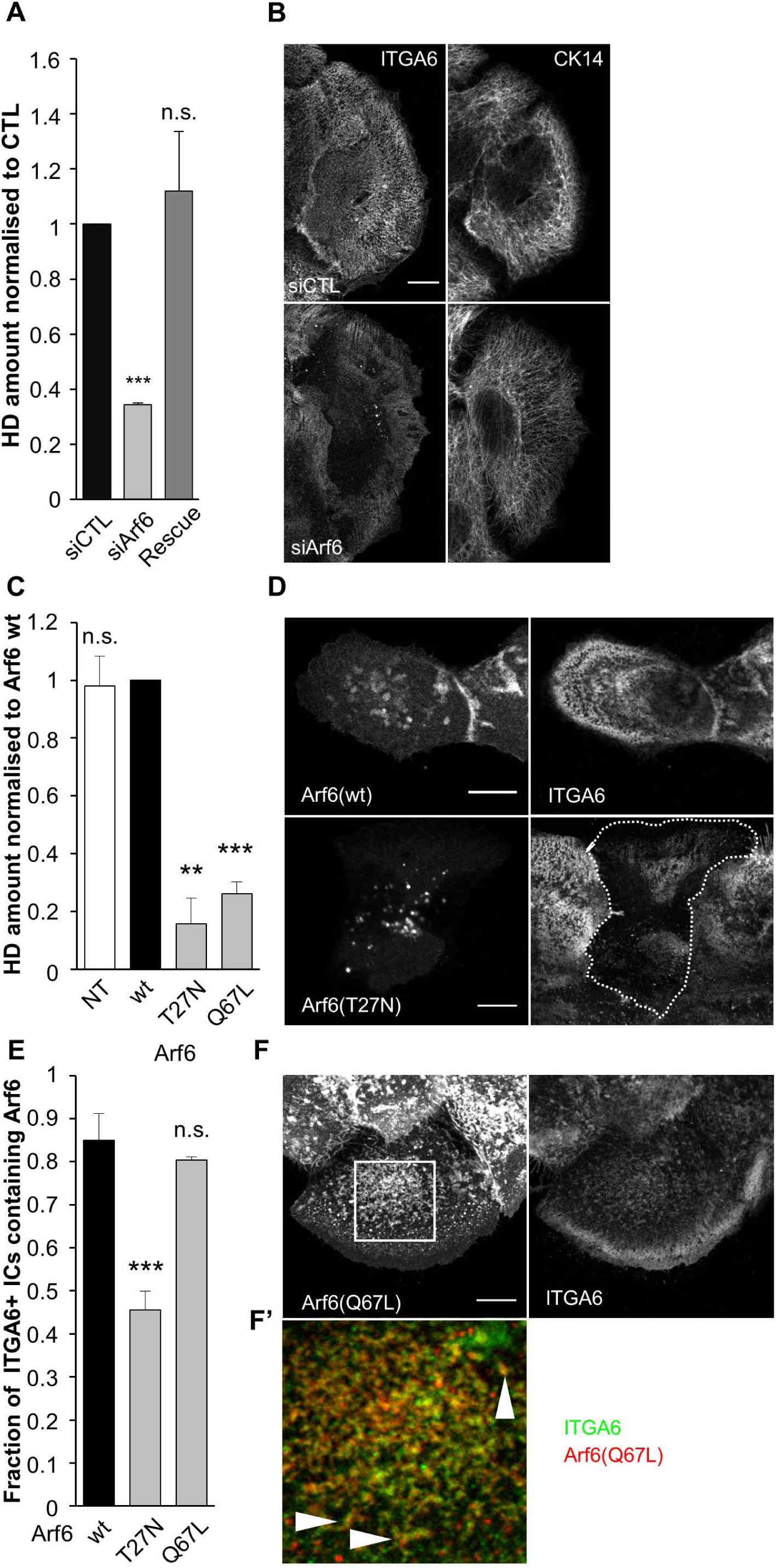
Arf6 is essential for proper HD organisation. (A) Quantification of the amount of HDs on the basal membrane of cells transfected with control siRNA (siCTL), Arf6 siRNAs (siArf6), or Arf6 siRNAs plus a plasmid encoding wild-type RFP-Arf6 plasmid mutated to be Arf6 siRNA-resistant (Rescue). (B) Confocal images of the basal membrane of HaCaT cells transfected with control or Arf6 siRNAs then stained for ITGA6 (left panels) and keratin 14 (CK14, right panels). Scale bar=10 μm. (C) Quantification of the amount of HDs on the basal membrane of cells expressing mutant forms of Arf6. (D) Confocal images of the basal membrane of cells expressing EGFP-Arf6(wt) or (T27N) immunostained for ITGA6 and CK14. Scale bar=10 μm. (E) Quantification of the presence of Arf6 in ITGA6-enriched intracellular compartments in cells expressing mutant forms of Arf6. wt=45, T27N=44, Q67L=46 cells. (F) Confocal images of the basal membrane of cells expressing EGFP-Arf6(Q67L) immunostained for ITGA6 and CK14. Scale bar=10 μm. (F’) Magnification of the boxed region in (F; magnification x3). Arrowheads highlight the colocalisation of Arf6 (red) and ITGA6 (green) at the basal plasma membrane.

Cav1 depletion also reduced HDs at the basal plasma membrane, but did not strongly affect keratins (Fig. 6A-B). Cav1 is a major caveolae component but also has a caveolae-independent membrane scaffolding function (Lajoie et al., 2009). To distinguish between both functions, we specifically affected caveolae by depleting Cavin1, another key caveolae component (Gambin et al., 2014). Cavin1 depletion reduced HDs at the basal plasma membrane but did not affect keratin organization to the same magnitude as Cav1 depletion (Fig. 6C-D). We also confirmed by transmission electron microscopy that Cav1- and Cavin1-depleted cells had significantly less caveolae at the basal plasma membrane (Fig S4A). This suggests that Cav1 regulation of HDs depends on caveolae. In other cell types, Cav1 function is regulated by Src-mediated Y14 phosphorylation (Parton and del Pozo, 2013). To test whether Cav1 phosphorylation alters its function in HD biogenesis, we expressed a phospho-mimetic Cav1(Y14D) or a non-phosphorylatable Cav1(Y14F) mutant form. Both forms significantly decreased HDs at the basal plasma membrane, with Cav1(Y14D) having stronger effects, while we did not measure significant difference between Cav1 wt-expressing cells and non-transfected cells (Fig. 6C-D).

**Figure 6.**
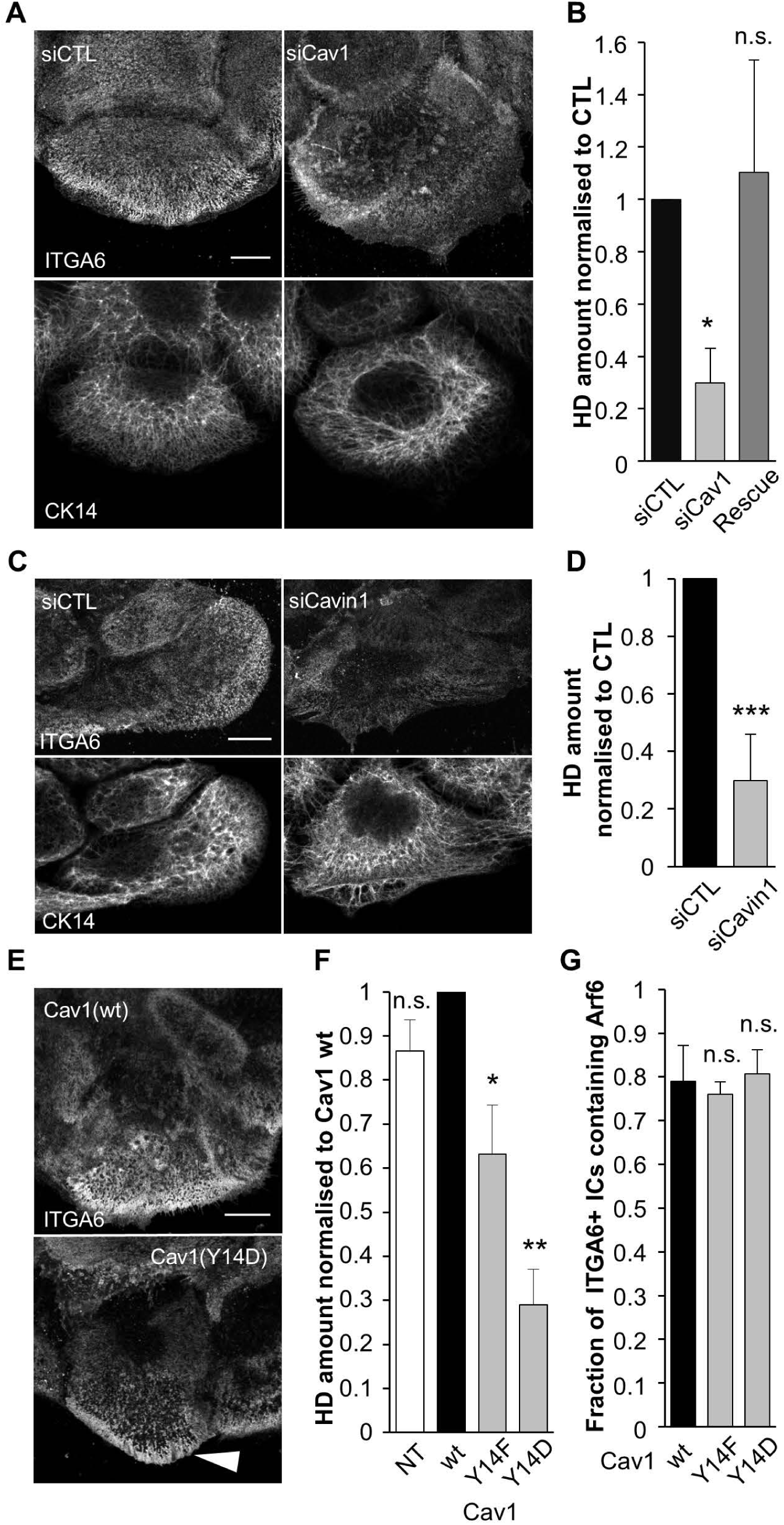
Cav1 and caveolae are essential for proper HD organisation. (A-D) Confocal images of the basal membrane of cells transfected with control, Cav1 siRNAs (A, B) or Cavin1 siRNAs (C, D) then immunostained for ITGA6 and CK14 (A, C); (B, D) quantification of the amount of HDs on the basal membrane of cells. The Rescue column (B) corresponds to cells transfected with Cav1 siRNAs plus a plasmid encoding wild-type RFP-Cav1 mutated to be Cav1 siRNA-resistant. Scale bar=10 μm. (E) Confocal images of the basal membrane of cells expressing wt or mutant forms of RFP-Cav1 immunostained for ITGA6 and CK14. Scale bar=10 μm. The white arrowheads show integrin accumulation at the edge of the cell in Cav1(Y14D) expressing cells. (F) Quantification of the amount of HDs on the basal membrane of cells expressing Cav1 mutant forms. (G) Quantification of the presence of Arf6 in ITGA6-enriched intracellular compartments in cells expressing mutant forms of Cav1. N: wt=27, Y14F=38, Y14D=33 cells.

Collectively, our data strongly suggest that Arf6 and Cav1 colocalize with ITGA6 at the basal plasma membrane as well as in internal compartments, and that HD formation requires Arf6 and caveolae.

### Arf6 and caveolae are essential for HD integrin dynamics

To further define the role of Arf6 and Cav1, and define whether they control HD formation by regulating α6β4 integrin trafficking, we examined their function in integrin turnover and in integrin endocytosis assays. First, we used confocal video-microscopy on cells expressing EGFP-β4 integrin (ITGB4) and mCherry-Arf6(wt) or RFP-Cav1(wt) to track Arf6-positive or Cav1-positive vesicles containing ITGB4. We saw that HD integrins were co-transported with Arf6 and Cav1 in such vesicles (Fig. 7A-B, Videos 1-2). Second, we measured integrin dynamics using FRAP in HDs. In wild-type keratinocytes expressing EGFP-ITGB4, the integrin mobile fraction was approximately 50% (Fig. 7C-D). In keratinocytes expressing the dominant-negative Arf6(T27N) or the phospho-mimetic Cav1(Y14D) mutant, the mobile fraction dropped to 36±7% or 34±2%, respectively, without any significant change in the half-time recovery (τ_1/2_, Fig. 7C-D). The constitutively-active Arf6(Q67L) and the non-phosphorylatable Cav1(Y14F) mutants did not affect ITGB4 dynamics (Fig. 7C-D). Hence, affecting Arf6 activity or mimicking Cav1 phosphorylation compromise the effective turnover of HD integrins.

**Figure 7.**
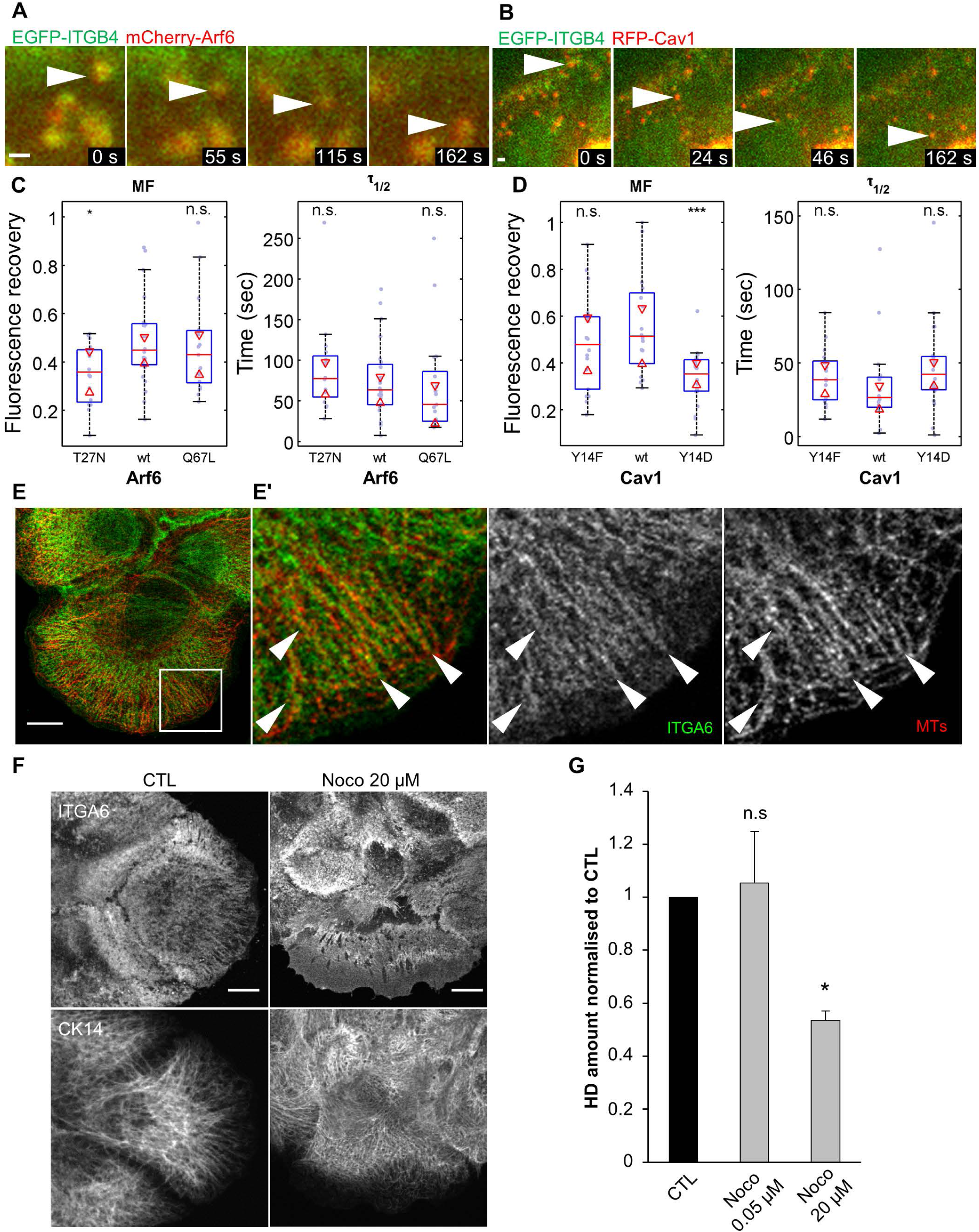
Arf6/microtubules-dependent traffic and Cav1 regulate HD integrin dynamics. Time lapse acquisition of EGFP-ITGB4 (green) and (A) mCherry-Arf6(wt) (red) extracted from video 1, or (B) RFP-Cav1(wt) (red) extracted from video 2. Time is shown in seconds. (C-D) FRAP analysis of EGFP-ITGB4 in HDs of cells expressing m-Cherry-Arf6 (C) or RFP-Cav1 (D) constructs. Left panel, mobile fraction (MF); right panel half-time recovery (τ_1/2_). (E) Confocal plane of the basal membrane of a cell stained for ITGA6 (green) and microtubules (MTs, red). (E’) Close-up of the squared region. Scale bar = 10 μm. (F) Cells were incubated with vehicle (CTL) or nocodazole (Noco) before fixation and immunostained for ITGA6 and CK14. Scale bar=10 μm. (G) Quantification of the area covered by HDs on the basal membrane of cells treated with vehicle or the MT depolymerizing drug nocodazole.

The directional motions of Arf6+/ITGB4+ and Cav1+/ITGB4+ vesicles suggests that they might move along microtubule (MT) tracks (Soldati and Schliwa, 2006). Consistent with this notion, ITGA6 often formed rod-like structures running parallel to MTs at the cell edge, presumably where MT tips are located (Fig. 7E-E’). Furthermore, MT depolymerization with high doses of nocodazole (20 μM) severely affected HDs at the basal plasma membrane with a diffuse localization of ITGA6; doses that inhibit MT dynamics without compromising their structure (0.05 μM) had no effect (Fig. 7F-G) (Liao et al., 1995; Vasquez et al., 1997). This is also supported by our recent data showing that MTs are required to address the adhesion receptor LET-805 to CeHDs in *C. elegans* epidermal cells (Quintin et al., 2016).

To define whether integrin turnover involves its trafficking between recycling vesicles and the plasma membrane, we next monitored the uptake of GoH3 antibodies specific for ITGA6. In control cells, we observed that ITGA6-bound GoH3 antibodies were localized in ITGB4+ ICs. In Arf6-, GIT1- or βPIX-depleted cells, ITGA6-GoH3 uptake was significantly reduced (Fig. 8A-B). By contrast, Cav1 or Cavin1 depletion had no effect on ITGA6 endocytosis (Fig. 8B). We confirmed these observations by quantifying biochemically the role of Arf6 and Cav1 on α6β4 integrin internalization. Arf6 depletion significantly decreased the endocytosis of both ITGA6 and ITGB4, suggesting that Arf6 is a key regulator of HD integrin turnover. Once again, Cav1 depletion did not significantly affect ITGA6 endocytosis, and only mildly reduced that of ITGB4, suggesting it may only affect ITGB4 subunits that are not incorporated within HDs (Fig. 8C), as previously reported (Seltmann et al., 2015). Finally we found that inhibition of dynamin activity using Dynasore did not affect HDs (Fig. 8D), nor ITGA6 uptake (Fig. 8E), while it prevented the uptake of EGF-bound EGFR (Fig. 8G), in contrast to what has been found for FA integrin recycling (Chao and Kunz, 2009; Ezratty et al., 2009) or caveolae internalization (Bass et al., 2011; Upla et al., 2004). This was further supported by the observation that dynasore did not affect HD integrin turnover measured by FRAP (Fig. 8E). These data support the idea that HD integrin turnover is regulated through the Arf6-dependent dynamin-independent pathway.

**Figure 8.**
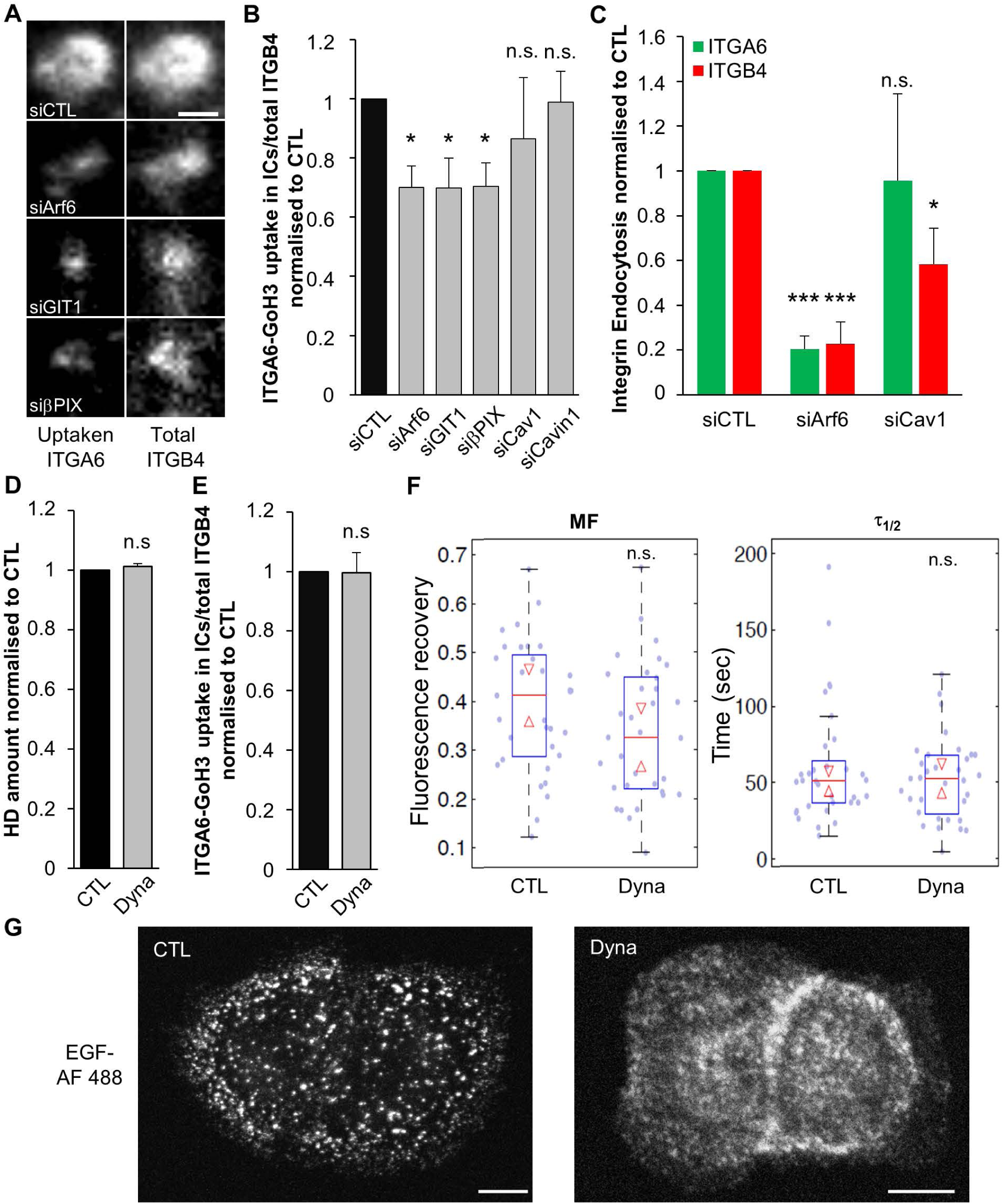
HD integrin endocytosis requires Arf6. (A-B) Antibody uptake assay after incubating HaCaT cells with the monoclonal antibody GoH3 (anti-ITGA6) on ice and shifting cells to 37°C for 1 hour. (A) Representative confocal images of ICs +1 μm above HDs in siRNA-transfected cells showing uptaken GoH3 bound to ITGA6 (left panel) and total ITGB4 (right panel). Scale bar = 1 μm. (B) Ratio of internalised GoH3-ITGA6/total ITGB4 measured in several ICs of individual cells for each condition following siRNA transfection. siCTL=30/262, siArf6=27/185, siGIT1 =29/130, siβPIX=25/145, siCav1=30/218, siCavin 1=25/196 cells/ICs. (C) Amount of endocytosed ITGA6 and ITGB4 integrins, following cell surface biotinylation at 4°C followed by internalisation at 37°C for 30 min. The amount of biotinylated α6 and β4 integrins was measured by capture ELISA with peroxidase-conjugated streptavidin, corrected for the total amount of ITGA6 or ITGB4 measured by western-blotting; ratios are expressed relative to control cells. (D) Quantification of the area covered by HDs at the basal membrane of cells treated with vehicle or the dynamin inhibitor Dynasore (Dyna). (E) Ratio of internalised GoH3-ITGA6/total ITGB4 measured in several ICs of individual cells for each condition following treatement with vehicle or Dynasore. CTL=18/105, Dyna=18/131 cells/ICs. (F) FRAP analysis of EGFP-ITGB4 in HDs of cells treated with vehicle or the dynamin inhibitor Dynasore. Left panel, mobile fraction (MF); right panel half-time recovery (τ_1/2_). (G) Z-projection of a confocal stack showing HaCaT cells fixed after 10 min incubation with EGF-AlexaFluor 488 in the presence of vehicle or dynasore. Scale bar = 10 μm. PDMS

Depending on the cell type, Cav1 Y14 phosphorylation either drives dynamin-dependent caveolae internalization (del Pozo et al., 2005), which does not seem to be the case in our model (see above) or promotes caveolar coat reorganization and caveolae swelling (Zimnicka et al., 2016). We examined integrin localization in Cav1(Y14D)-expressing cells and found that in 37/59 cells the HD remnants were present at the cell edge (Fig. 6C), potentially coinciding with MT tips. One interpretation, consistent with the decrease in ITGB4 mobile fraction and the effect of Cav1-Y14 phosphorylation, could be that Cav1(Y14D) prevents HD integrin turnover. Alternatively, it could slow down integrin recycling and induce their accumulation at the cell edge, thereby reducing the total amount of HDs during cell spreading.

We conclude that α6β4 integrin recycling is regulated though the Arf6-dependent dynamin-independent pathway and that functional caveolae are required for α6β4 integrin stabilization at the basal plasma membrane. These two steps are critical for HD formation and maintenance.

### Arf6 and GIT1/βPIX regulate Cav1 targeting to the basal plasma membrane

The data described above suggest that Arf6 acts mainly along a trafficking route, whereas Cav1 acts mainly at the basal plasma membrane. To test this prediction by other means, we examined the mutual requirement for basal plasma membrane and IC localization of Arf6, GIT1, βPIX and Cav1. Using siRNA-mediated depletion or the transfection of mutant forms we examined the effects on the other players. Our data strongly suggest that GIT1 and βPIX basal plasma membrane localization depends on normal Arf6 activity, and that all three factors are required to transport Cav1 to the basal plasma membrane (Fig. S4B-E, Video 3). It confirms previous reports showing that GIT1 and βPIX shuttle between endosomes and the membrane in an Arf6-dependent manner (Di Cesare et al., 2000; Matafora et al., 2001; Osmani et al., 2010; Valdes et al., 2011). Importantly, we found that GIT1 and βPIX did not affect Cav1 phosphorylation despite their requirement for its localization (Fig. S4F). Conversely, we found that expressing Cav1(Y14D) did not reduce the proportion of ITGA6-positive/Arf6-positive ICs compared to control cells (Fig. 6E). Finally, we found that Rac1 and PAK1 act at a different step, since PAK1-or Rac1-depleted cells had normal levels of Cav1 at the basal plasma membrane (Fig. S4D-E). Collectively, our data suggest that Cav1 acts downstream of Arf6 and GIT1/βPIX to favor HD formation and strengthens the notion that it acts at the basal plasma membrane.

### HD remodeling in response to mechanical stress depends on Cav1 and Arf6

The results described so far identify a pathway promoting HD biogenesis. It might prove particularly necessary when tissues experience higher mechanical load such as during growth or wound healing. To directly assess whether HDs respond to mechanical cues, as in *C. elegans* (Zhang et al., 2011), we used a home-built stretching device compatible with confocal live imaging to apply a constant uniaxial 10% stretch on the cell substrate (Fig. 9A). Immediately after stretch, HDs were passively stretched: the cell area increased by 10%, while HD density (HD integrin concentration at the membrane) decreased by 10% (Fig. 9B). In parallel, immediately after stretching we observed a ∼40% decrease of Cav1(wt) at the basal plasma membrane. Following this initial response, the HD amount increased by 20% over 30 min and maintained constant density, suggesting a mechanically-induced recruitment of HD integrins (Fig. 9B,D-G, Video 4). Under strain, the increased HD amount correlated with a decrease of Cav1(wt) at the basal plasma membrane where HDs appeared (Fig. 9C-F,I, Video 4). The analysis of the HD mechanoresponse over time showed a linear increase in HD amount, indicating an active process with a median speed of 1.5±0.6 μm^2^/min. We also measured a density enrichment of integrins in HDs of 2.6±0.7 compared to integrins diffusing in the membrane close to the surface. These values are consistent with a diffusion mechanism for integrin recruitment when the membrane is flattened. Interestingly, these values are consistent with the reported measures of ∼1 μm^2^/min for β1 integrin diffusion and a 2-fold density enrichment in FAs compared to diffusing integrins close to the surface (Rossier et al., 2012). This is also consistent with the growth mechanism through membrane flattening and diffusion, first reported for cadherin-based adhesions (Delanoë-Ayari et al., 2004). However, in contrast to FAs (Riveline et al., 2001) and adherens junctions (AJs) which grow along the direction of stretching (Brevier et al., 2007; Brevier et al., 2008), HD amount increased in all directions (Fig. 9H).

**Figure 9.**
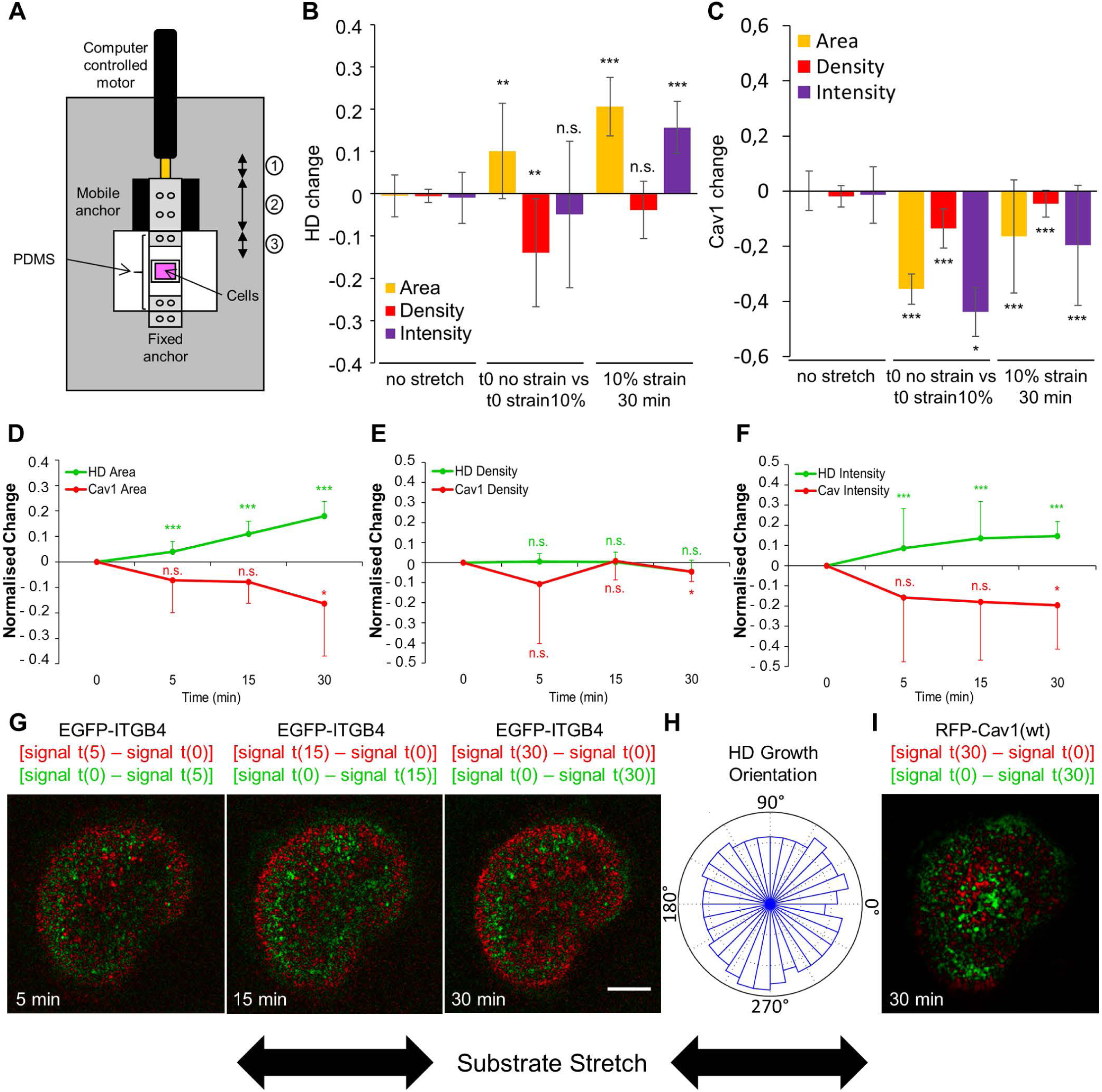
HDs respond to external mechanical stimuli. (A) Scheme of the custom-built stretcher used. The motor motion (1) induces the displacement of the mobile anchor (2) driving the stretching of the PDMS membrane (3). (B-C) Quantification at the basal plasma membrane of the area (yellow), density (red) and total intensity (purple) changes of (B) EGFP-ITGB4 or (C) RFP-Cav1(wt) without strain (left), before and immediately after a 10% uniaxial strain (middle), or after 30 min of a 10% uniaxial strain (right). (D-F) Quantification of the area (D), total intensity (E) and density (F) of HDs (green) and Cav1 (red) in cells expressing EGFP-ITGB4 and RFP-Cav1(wt) after 5, 15 and 30 min of stretch with a 10% uniaxial strain. All values are normalised to time 0. (G,I) Confocal images of a cell expressing EGFP-ITGB4 (G) and RFP-Cav1 (I) under a 10% uniaxial strain after 5, 15 and 30 min. The green channel shows what disappears by subtracting from the signal in the first frame t(0) the signal at time t (5, 15 or 30 min): signal t(0) – signal t(x); the red channel shows what appears by subtracting from the signal at time t (5, 15 or 30 min) the signal in the first frame t(0): signal t(x) – signal t(0). Scale bar = 10 μm. (H) Quantification of the orientation of HD growth after 30 min of strain. N = 12 cells. Images are related to video 4.

Since Arf6 and caveolae components were required for HD formation and integrin turnover, we tested their role in the mechanoresponse of HDs. In Cav1(Y14D)- or Arf6(T27N)-expressing cells, HDs were unable to respond to mechanical strain and rather disappeared, while Cav1(Y14D) was not removed (Fig. 10A-D). Recent results have suggested that caveolae can act as a mechanical buffer by flattening in response to the change in membrane tension, whether induced by cell stretching or a hypotonic shock (Sinha et al., 2011). To test whether HD growth was connected to a similar mechanically-induced caveolar flattening, we also used osmotic swelling as an alternate mechanical cue. We observed that a 10-fold decrease in osmolarity led to a 2-fold increase in HD area correlated with a 2-fold decrease in caveolae at the basal plasma membrane (Fig. 10E-G). This effect was dependent on caveolae integrity, since HDs did not respond to mechanical input in Cavin1-depleted cells (Fig. 10E). Finally to assess whether mechanically-induced HD growth relied on the Arf6-dependent recycling of HD integrins from the ICs, we measured the mobile fraction of EGFP-ITGB4 either in isotonic or hypotonic conditions. Focusing specifically on ICs, we observed a 69% recovery of EGFP-ITGB4 in cells grown in isotonic medium. By contrast, we failed to observe any recovery after a 10 min osmotic shock (Fig. 10H). These results strongly support the notion that mechanically driven HD growth requires active shuttling of the pool of free HD integrin available in the ICs towards the plasma membrane.

**Figure 10.**
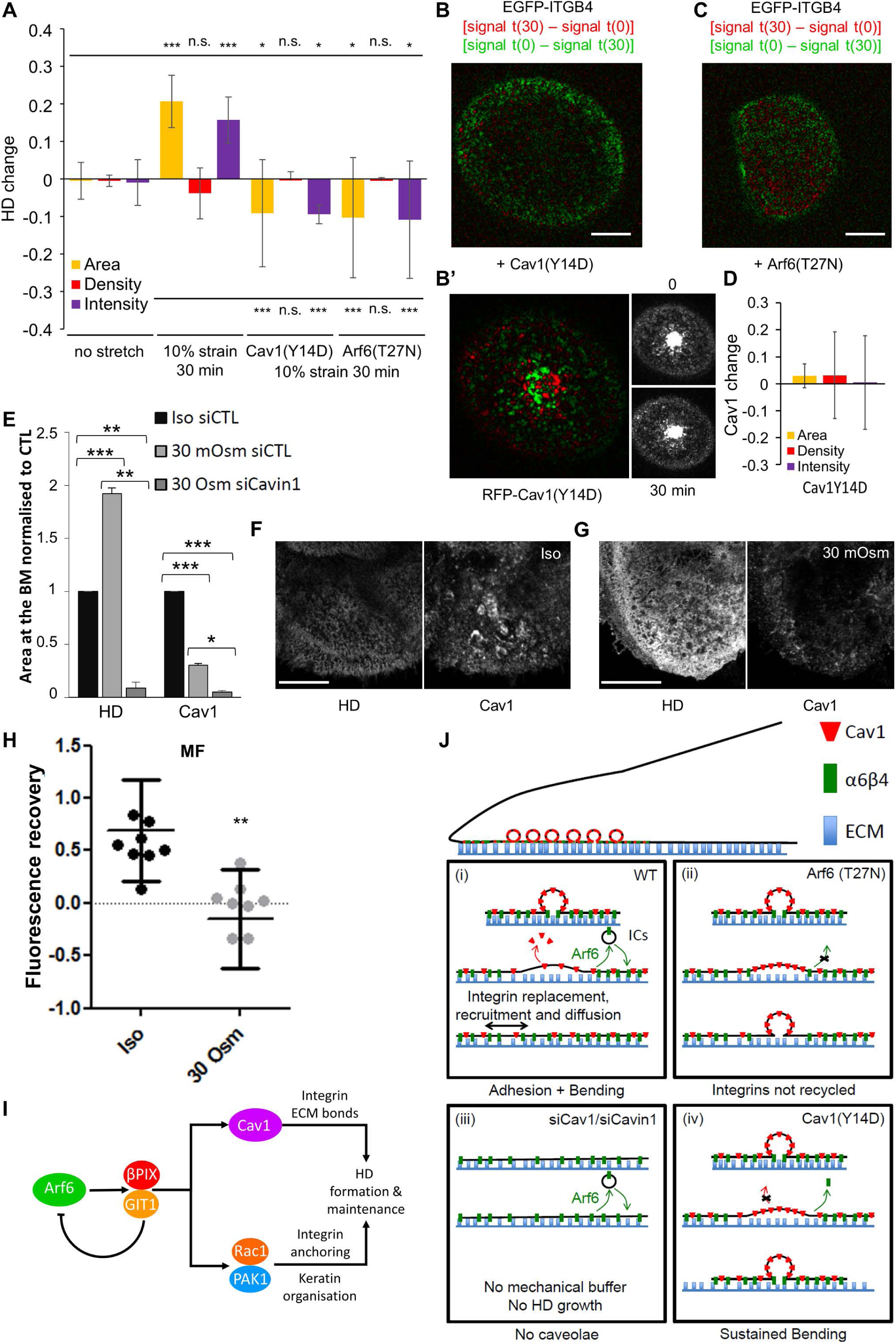
Cav1 and Arf6 are required for proper HD mechanoresponse to external forces. (A) Quantification of the area (yellow), density (red) and total intensity (purple) changes of HDs after 30 min without stretch in cells expressing EGFP-ITGB4 (N=20) or after 30 min of a 10% uniaxial stretch in cells expressing EGFP-ITGB4 and RFP-Cav1(wt) (N=12), RFP-Cav1(Y14D) (N=14) or mCherry-Arf6(T27N) (N=11). (B-C) Representative confocal images of the basal membrane of a cell submitted to a 10% strain during 30 min expressing EGFP-ITGB4 and (B-B’) RFP-Cav1(Y14D) or (C) Arf6(T27N); image representation as in Fig. 9. (D) Quantification of the area (yellow), density (red) and total intensity (purple) changes of Cav1(Y14D) after 30 min of a 10% uniaxial stretch in RFP-Cav1(Y14D) (N=14, related to Fig. 10B’). (E) Quantification of the amount of HDs and Cav1 at the basal membrane of cells in isotonic medium (Iso: 300 mOsm) or after a 10 min hypotonic shock (30 mOsm) in control cells or cells transfected with siCavin1 siRNAs to remove caveolae. (F-G) Confocal images of the basal membrane of cells stained for ITGA6 and Cav1 in (F) isotonic medium (300 mOsm) or (G) after a 10 min hypotonic shock (30 mOsm). Brightness and contrasts are unmodified. Scale bar=10 μm. (H) Mobile fraction (MF) of EGFP-ITGB4 in the ICs of cells incubated with isotonic medium (300 mOsm) or after a 10 min hypotonic shock (30 mOsm). (I-J) Proposed mechanism for the role of Arf6, GIT1 /βPIX, PAK1/Rac1 and caveolae components in HD remodeling (I) and for HD mechanoresponse (J). See discussion for details.

Altogether, our data strongly suggest that we have identified a novel trafficking-dependent pathway involved in mechanoresponse at HDs relying on Arf6 and Cav1.

## Discussion

In this study, we characterize a pathway mediating HD remodeling in keratinocytes during cell spreading, and in response to mechanical tension. From a molecular standpoint, this pathway features a central role for Arf6 and GIT1/βPIX in shuttling HD integrins between presumptive recycling endosomes and the basal plasma membrane. From a mechanical standpoint, caveolae play a key role by allowing HDs to remodel and α6β4 integrin to bind the ECM, probably as further discussed below by flattening under stress. Finally, Rac1 and PAK1, which are probably co-transported with integrins, mediate the reorganization of the keratin network and reinforce HDs at the basal plasma membrane at a later step (Fig. 10I). Our analysis of HD mechanoresponse reveals both similarities and differences with FA mechanoresponse to mechanical stress.

Three lines of evidence support the notion that Arf6, a regulator of membrane trafficking (Donaldson and Jackson, 2011), together with GIT1 and βPIX act upstream in HD formation. First, HD integrins are shuttled between large compartments containing Arf6/GIT1/βPIX and the basal plasma membrane depending on Arf6 (Fig. 10H). Second, Arf6 is required to target GIT1 and βPIX to the basal plasma membrane (Frank and Hansen, 2008), and conversely the Arf6-GAP activity of GIT1 (Meyer et al., 2006) is required for HD formation, suggesting their mutual dependence in transport. Finally, Arf6 overactivation induced an abnormal accumulation of integrins at the basal plasma membrane. We thus suggest that Arf6 regulates the endocytosis and the recycling of HD integrins, and that HD formation requires a dynamic shuttling of α6β4 integrins.

We argue that Cav1, and presumably also Cavin1, act downstream of Arf6/GIT1/βPIX, since the latter were needed for Cav1 basal plasma membrane localization. We further argue that Cav1 and Cavin1 are required for HD integrin turnover through a mechanism that does not involve Cav1-mediated endocytosis. Indeed, depleting Cavin1, which has not been associated with caveolae-mediated endocytosis, affects integrin turnover in mechanical assays (see below), but not its uptake by endocytosis. Moreover, dynamin inhibition had no effect on HDs, nor on integrin uptake, suggesting that HD integrin endocytosis occurs through an Arf6-dependent recycling pathway (Doherty and McMahon, 2009), rather than through dynamin-dependent caveolae internalization as reported in some cells (del Pozo et al., 2005). Lastly, expressing the phospho-mimetic mutant Cav1(Y14D) reduced HD integrin mobility and induced their abnormal accumulation at the cell edge where MT tips are located. This abnormal accumulation could indicate that caveolae are indirectly required to enable HD integrin trafficking through an Arf6-dependent pathway. Below, we propose a model to reconcile these data and account for the role of caveolae and Arf6 in HD remodeling.

One of our key findings is that HDs are mechanoresponsive since they grow under external forces. We propose that the mechanism involved in this response relies on the mechanical stress buffer function of caveolae (Sinha et al., 2011), and on Arf6-dependent integrin recycling. As previously reported for FAs (Balaban et al., 2001; Galbraith et al., 2002; Moore et al., 2010; Parsons et al., 2010; Riveline et al., 2001) and adherens junctions (Delanoë-Ayari et al., 2004; Brevier et al., 2007; Brevier et al., 2008), we found that HD intensity and area, but not density, increased under external forces, strongly suggesting that integrin recruitment depends on mechanical tension. However, whereas FA/AJ growth directions are set by actomyosin force orientation, HD integrin recruitment was not directional. Two arguments suggest that the non-directionality of growth results from the isotropic flattening of caveolae. First, stretching was associated with an immediate decrease in integrin density (integrin concentration in HDs), suggesting that new membrane devoid of integrin appears at the basal plasma membrane in close vicinity of the ECM, and with Cav1 disappearance, indicating its unbinding from caveolae during flattening. Second, caveolae have a mostly “spherical” symmetry, hence their flattening should be isotropic.

Biophysical studies have suggested that the size specificity of caveolae results from a competition between the bending energy induced by Cav1 and the large surface tension of the plasma membrane (Sens and Turner, 2004, 2006). We propose that, upon stretching, caveolae are flattened and the new membrane facing the ECM can either return to its caveolar Ω-shape because the bending energy is favored, or remain flat if α6β4 integrins invade the newly available membrane by diffusion from basal plasma membrane or transport from RE and form new bonds thereby stabilizing new HD adhesions (Fig. 10J). This mechanism is encoded in a competition between caveolar flattening and integrin/ECM adhesion energies. We further propose that HDs serve as pinning points in the vicinity of caveolae that must be removed by Arf6-dependent HD recycling (i) to release adhesions and allow spreading; (ii) to promote the formation of HD complexes at new adhesion sites (Fig. 10Ji). Compromising Arf6 activity prevents such a recycling, thereby yielding to membrane bending and caveolae re-formation (Fig 10Jii). Likewise, we suggest that compromising with Cav1 or Cavin1 function (Figs 6A-D, 10E) denies cells from any mechanical buffer and prevents basal plasma membrane reorganization (Fig. 10Jiii). Finally, expressing the phosphomimetic Cav1(Y14D) form prevents Cav1 unbinding from caveolae, thus energetically favoring their return to the caveolar Ω-shape (Fig. 10Jiv). The absence of mechanically-induced flattening in Cav1(Y14D)-expressing cells argues that Cav1 phosphorylation at Y14 is critical for bending.

Experimental considerations can also account for why growth is restricted to the cell edge over a region of ∼1 μm, where microtubule tips are also found, highlighting a transport-mediated local regulation of adhesion (Fig. 7E-E’). First, it has been proposed that the local competition between spreading of a spherical cell and adhesion triggers an increase in membrane line tension at the very edge (Bar-Ziv et al., 1999). Moreover, we previously showed that the cell cortex locally senses the force over ∼1 μm distance (Delanoë-Ayari et al., 2004), which is consistent with the HD growth from the cell border. Cav1(Y14D) induced accumulation of HD integrins at the very edge (Fig. 6C), suggesting the lack of caveolae mechanoresponse, prevents HD integrin reorganization and consequently HD growth. Finally, a recent super-resolution mapping of HD components suggests that HDs forming at the edge of the cell are lacking BP180 and BP230 (so-called type II HDs), and might thus be easier to reorganize than type I HDs including BP180 (so-called type I HDs) (Nahidiazar et al., 2015). Altogether, the HD mechanical response appears to be mediated by the cortex in response to the local force.

Interestingly, while we emulated the strategy used in *C. elegans* to identify factors acting in HD remodeling (Zahreddine et al., 2010; Zhang et al., 2011), the precise mechanosensitive response observed in vertebrates appears different, featuring a prominent role for integrin trafficking. Two important differences are that *C. elegans* does not form caveolae (Kirkham et al., 2008), most likely due to the absence of a Cavin homologue in its genome, nor organizes its CeHDs with integrins (Osmani and Labouesse, 2015).

Finally, we suggest that Rac1 and its kinase effector PAK1 act downstream of Arf6/GIT1/βPIX, and in parallel to Cav1, in HD remodeling. Indeed, although Rac1 and PAK1 were present in the Arf6/HD integrin-enriched compartment, they were not required to localize Cav1. Yet the βPIX RacGEF activity and βPIX-PAK1 interaction affected HDs but not Cav1 localization. Instead, PAK1 might control keratin organization through phosphorylation, akin to what happens in *C. elegans* (Zhang et al., 2011) and supported by PAK1 direct phosphorylation of vimentin (Goto et al., 2002). Consistent with the report that keratinocytes lacking all keratins have reduced HDs (Seltmann et al., 2013), we suggest that PAK1-dependent keratin phosphorylation might promote the subsequent stabilization of new integrin clusters.

The phosphorylation of the ITGB4 cytoplasmic tail had been shown to be a major driver of HD disassembly (Frijns et al., 2012; Frijns et al., 2010; Germain et al., 2009; Rabinovitz et al., 2004; Wilhelmsen et al., 2007), but the pathways regulating their reassembly had remained elusive until the present work. Collectively, our data support a mechanism whereby in response to external mechanical cues caveolae flattening promotes integrin mobilization to reinforce HDs in an isotropic mode.

## Acknowledgements

We thank Nicolas Vitale for constructs and helpful discussions, the IGBMC Imaging Center for help with microscopy, Anita Eckly-Michel and Fabienne Proamer for their help with electron microscopy and Raghavan Thiagarajan for help in the micro-contact printing experiments. D.R. acknowledges Pierre Sens for discussions on the physical mechanism. We thank Bruno Klaholz and Leonid Andronov for granting access to the GSD microscope, which is supported by the French Infrastructure for Integrated Structural Biology (FRISBI) ANR-10-INSB-05-01, and Instruct as part of the European Strategy Forum on Research Infrastructures (ESFRI). We also thank Romeo Ricci, Nicolas Vitale, Patricia Simon-Assman, Sophie Quintin and Adèle de Arcangelis for critical reading of the manuscript. This work was supported by Agence Nationale de la Recherche and European Research Council grants to ML, by grants from ISIS, the ci-FRC of Strasbourg and Fondation Simone et Cino del Duca to DR, by institutional funds from INSERM and University to JGG, by the grant ANR-10-LABX-0030-INRT, a French State fund managed by the Agence Nationale de la Recherche under the frame programme Investissements d’Avenir labelled ANR-10-IDEX-0002-02 to the IGBMC and by institutional funds from the CNRS, INSERM, University of Strasbourg.

The authors declare no competing financial interests.

## Author contributions

ML and EGL proposed the project. ML and NO designed the experiments; NO carried all cell experiments. JP designed image analysis programs and performed the analysis. JC and DR conceived the cell stretcher and the early spreading/pattern experiments and designed and developed the physical analysis. JC helped with cell stretching experiments. NF prepared electron microscopy samples, acquired and quantified data. JGG helped with the design and analysis of EM, caveolae/Cav1 experiments and provided Cav1 constructs. NO, JP, JC, DR and ML analyzed the data. NO and ML wrote the manuscript with inputs from all authors.

## Materials & methods

### Plasmids and siRNAs

pEGFP-H1B was a gift from Edgar Gomes (IMM, Lisbon, Portugal). pcDNA3-ITGB4-EGFP was a gift from Arnoud Sonnenberg (NKI, Amsterdam, The Netherlands). All βPIX, PAK1, GIT1 and Rac1 as well as pEGFP-Arf6 constructs were gifts from Nicolas Vitale (INCI, Strasbourg, France). pmCherry-Arf6 constructs were generated by subcloning pEGFP-Arf6 constructs into a pmCherry-N1 vector. pLKO.1-puro-shPlectin (TRCN0000082813) targeting human plectin was purchased from Sigma. All siRNAs are ON-TARGETplus SMARTpools and were purchased from Dharmacon. We confirmed that the siRNAs specifically depleted at least 75% of each targeted mRNA (Fig. S5). SMART pool siRNA resistant human Arf6 and Cav1 cDNAs with silent mutations on siRNA targeted regions were generated and cloned in pD673-CRc containing mCayenneRFP in C-terminal position by DNA2.0.

### Antibodies

Antibodies against ITGA6 (rat monoclonal GoH3), ITGB4 (mouse monoclonal 7/CD104), Cav1 (mouse monoclonal 2297/Caveolin1 and polyclonal rabbit), Rab11 (mouse monoclonal 47/Rab11) and EEA1 (mouse monoclonal 14/EEA1) were purchased from BD Transduction Laboratories; against CK14 (rabbit polyclonal) from Convance; against Plectin (polyclonal Guinea Pig) from Progen; against βPIX, PAK1 and ITGB1 (polyclonal rabbit) from Millipore; against Arf6 (rabbit polyclonal) from AbCam; against pY14-Cav1 (polyclonal rabbit) from Cell Signaling; against pY397-FAK (polyclonal rabbit) from Invitrogen; against α-tubulin (monoclonal mouse DM1A), FLAG (monoclonal mouse M2 and rabbit polyclonal) and HA (rabbit polyclonal) from Sigma; against the HA tag (monoclonal rat 3F10) from Roche the antibody against the Myc tag (monoclonal mouse 9E10) was made in-house (IGBMC, Strasbourg). All secondary antibodies used in immunofluorescence stainings are coupled to Alexa Fluor 488, 594 or 647 dyes and were purchased from Molecular Probes.

### Drugs

FAK inhibitor PF-228 was purchased from Millipore and incubated with cells at 1 μM during 2h before fixation. Nocodazole and Dynasore were purchased from Sigma and incubated with cells at 0.05 (low concentration) and 20 (high concentration), and 80 μM, respectively, during 1h prior to fixation.

### Cell culture

HaCaT cells were obtained from the DKFZ (Heidelberg, Germany) and were cultured in DMEM 1 g/L glucose (Invitrogen) supplemented with 10% fetal calf serum (PAN Biotech) and 10 μg/ml gentamicine (Kalys). Stable GFP-H1B, shPlectin and shPlectin/GFP-H1B cell lines were respectively selected and maintained with 500 μg/ml geneticin (Invitrogen), 10 μg/ml puromycin (Sigma), 10 μg/ml puromycin and 500 μg/ml geneticin. For live imaging experiments, the medium was changed just before mounting cells on microscope slides with DMEM 1 g/L glucose supplemented with 10% foetal calf serum and 10 μg/ml gentamicine without phenol red and supplemented with 20 mM Hepes. For micro-contact printing experiments, poly-lysine circular motifs of 30μm in diameter were prepared along the protocol reported previously (Caballero et al., 2014). Cells were left to spread on micropatterns for 1h in medium without serum to reduce ECM deposition outside the micropatterns.

### Transfection

DNA plasmids were nucleofected in cells using Nucleofector device and Cell Line Nucleofector Kit V (Lonza) following the manufacturer’s instructions and experiments performed 24h later. siRNAs were transfected into cells with Lipofectamine RNAiMAX (Invitrogen) following the manufacturer’s instructions, and experiments were performed 72 h later.

### siRNA screen

GFP-H1B and GFP-H1B/shPlectin stable cell lines were seeded in 24-well plates and transfected with siRNAs for candidate genes. 72hours later, the confluent monolayer was wounded with a glass micropipette (approximately 300 μm wide). Cell migration was monitored using GFP/phase contrast time lapse video-microscopy over 24h at a rate 1 image/15 min using an inverted DMIRE2 microscope equipped with a HC PL FLUOTAR 10X 0.30 air objective and a 37°C incubation chamber (Leica). Movies were analyzed using Imaris (Bitplane). EGFP nuclei were detected and tracked over time with an autoregressive motion over a maximum distance 20 μm and no gap between frames were allowed. Cells moving less than half of the total migration time were excluded. Aberrant trajectories were manually excluded. For each siRNA a ratio (V_shPlec_)/(V_wt_), where V_shPlec_ and V_wt_ are the mean speed of shPlec and wt cells respectively, was calculated and normalized to the ratio for control siRNA. Genes potentially encoding HD remodeling factors should increase speed and result in a ratio

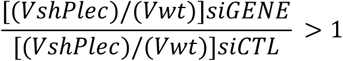

(see text and Fig. S1A for explanation).

### Immunofluorescence

Cells were fixed with paraformaldehyde (PFA) 4% 10 min at room temperature, permeabilized with PBS-0.2% Triton 10 min at room temperature. Aldehydes were reduced with PBS-NaBH_4_ for 10 min at room temperature. Primary antibodies diluted in PBS were incubated for 1h at room temperature and corresponding secondary antibodies diluted in PBS were incubated 30 min at room temperature.

### Confocal microscopy

All images from fixed samples were acquired with a TCS SP5 confocal microscope equipped with a HCX PL APO 63X NA 1.4 oil objective (Leica). All data sets were acquired in a row with the same settings.

### HD/Cav1/GIT1/βPIX image analysis

Single plane images corresponding to the plasma membrane were analyzed using the MetaMorph software (Molecular Devices). To quantify the area occupied by HDs, briefly, a region of interest (ROI) was manually drawn around each individual cell to isolate them. Control cells (control plasmids or siRNA) were used to determine a detection threshold using 4 distinct square ROI of 50 pixels placed randomly in the cell. The same threshold was applied on the rest of the sample (mutant constructs or siRNAs). For each cell, the areas of the whole cell and the thresholded signal at the basal membrane were measured to determine a ratio (thresholded signal area)/(cell area). Values were then normalized to values for control cells. Each experiment was reproduced at least 3 times independently with a total sample number of at least 30 cells.

### Keratin image analysis

We developed a new analysis method inspired from the physics of liquid crystal to quantify the organisation of the keratin cytoskeleton through nematic order (see supplementary fig. 1g for explanation). Single plane images corresponding to the plasma membrane were pre-processed in order to extract the bundles of filaments. First, the low frequencies (slow changes of intensity) were removed using a top-hat transform. Keratin bundles were then enhanced by computing the Laplacian operator of the images. Finally, skeletons of the filaments were extracted from the Laplacian image and the crossing pixels were removed.

Two measures were then computed on these skeletons: a neighborhood order coefficient (NOC) and a radial order coefficient (ROC) used to measure respectively the local pattern organization and the radial pattern organization of the filaments. For this purpose, the angular error 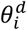 between the actual orientation of the skeleton at pixel *i* and the expected orientation given by a director *d* is computed. In the NOC case, the director is the average of orientations in a neighbourhood of size 21x21 pixels centred on pixel *i* whereas in the ROC case, the director is formed by the vector from the nuclear center to pixel *i*.

For pixel *i*, a local score *s_i_* is then obtained by taking:

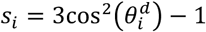
The higher the obtained value is, the more organised the cytoskeleton is at pixel *I* and conversely. Finally, the score of one image was obtained by averaging the scores of all considered pixels. All processing has been performed under *ImageJ* and using homemade macros.

### Anti-ITGA6 uptake

Cells were washed twice with ice-cold medium and incubated with anti-ITGA6 antibody (GoH3) at 1 μg/ml on ice for 30 min, before two additional washes with ice-cold medium. The internalization of ITGA6-bound antibodies was induced by adding prewarmed medium and incubating cells at 37°C for 1h. After fixation, permeabilization and PFA reduction, ITGA6-bound antibodies were revealed with an Alexa Fluor 488-conjugated anti-rat antibody and cells were immunostained for total ITGB4 with mouse monoclonal anti-ITGB4 and Alexa Fluor 647-conjugated anti-mouse antibody. Single confocal images at 1 μm above the basal membrane were acquired. ITGA6 internalization was measured using a custom-made macro in *ImageJ* software by drawing ROIs around individual ICs in each cell. The total intensities in the ITGA6 and ITGB4 channels were measured in each IC of individual cells for each condition. Internalized ITGA6 was normalized to total ITGB4 as the ratio [ITGA6 total intensity]/[ITGB4 total intensity] in each IC.

### Biochemical measure of integrins α6 and β4 internalization

HD integrin endocytosis was measured biochemically by adapting the protocol described in (Roberts et al., 2001). Cells were washed twice in cold PBS and their surface was labeled with 0.2 mg/ml NHS-SS-biotin in PBS for 30 min at 4°C. Cells were washed twice in cold PBS and surface proteins internalization was induced by adding pre-warmed medium and incubating cells at 37°C for 30 min. After removing the medium, cells were washed twice in cold PBS. Biotin was removed from proteins remaining at the cell surface by incubating cells in Tris 50mM pH 8.6 NaCl 100mM MesNa 20mM for 15 min at 4°C. MesNa was quenched by adding 20 mM of iodoacetamide for 10 min at 4°C. Cells were lysed in Tris 75 mM pH 8.6, NaCl 0.2 M, EDTA 7.5 mM, EGTA 7.5 mM, Igepal CA-630 0.75%, Triton X-100 1,5% and Complete Protease inhibitor cocktail (Roche). Lysates were fractioned with a 27G needle and centrifuged at 10000g 4°C for 10 min. The amount of biotinylated integrins α6 and β4 were measured by capture ELISA. Briefly, COSTAR 96 well High Bind EIA/RIA plates (Corning) were coated with each anti-integrin antibodies separately at 5 μg/ml in Na_2_CO_3_ 50 mM pH 9.6 overnight at 4°C. The plate was blocked with PBS-0.05% Tween-5% bovine serum albumin for 1h30 at room temperature. Integrins α6 and β4 were captured separately in duplicates by adding 50 μl lysates in each well for 2 h at RT under gentle agitation. Plates were extensively washed with PBS-0.05% Tween. Biotin was revealed by incubating wells with horseradish peroxidase conjugated-streptavidin (Pierce) in PBS-0.05% Tween-1% BSA for 1 h at RT. After washing, integrin biotinylation was measured by a chromogenic reaction with 3,3’,5,5’-Tetramethylbenzidine (TMB) ELISA substrate (Interchim). The amount of α6 and β4 integrins in each sample was measured by western-blotting for ELISA normalization.

### Super-resolution microscopy and image analysis

Samples were immunostained as mentioned above using Alexa Fluor 488 and 647-conjugated secondary antibodies in glass bottom dishes (μDish 35 mm high, ibidi) and set in a 10 mM MEA, 25 mM Hepes in PBS. All images were acquired with a Leica SR GSD TIRF microscope equipped with a HCX PL APO 100 X NA 1.47 oil objective and a 1.6X lens and a iXON ULTRA DU897V EM-CCD camera (Andor). Super-resolution images were reconstructed from GSD raw images using a non-parametric kernel smoothing custom-made method with *ImageJ* software. Colocalization was measured using MATLAB (Mathworks) with the Pearson’s correlation coefficient defined as

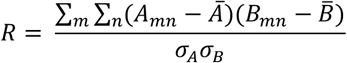
where 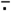 is the average and *σ*. the standard deviation. For βPIX and PAK1, the correlation was measured specifically in focal adhesions, isolated with a specific ROI around them, and in the rest of the cell. The results were normalized to the correlation value of the positive control ITGA6/ITGB4.

### Live imaging and FRAP experiments

Cells were seeded on glass bottom dishes (μDish 35mm high, ibidi). Time-lapse acquisitions were performed on cells expressing EGFP-ITGB4(wt) and RFP-Arf6 or Cav1(wt or mutants) with an inverted Zeiss Axio Observer Z1 microscope equipped with a PLAN APO 100X NA 1.4 oil objective (Zeiss), a Yokogawa CSU-X1 confocal head, an Evolve 512 EM-CCD camera (Photometrics), an iLAS FRAP illumination module (Roper Scientific) and a 37°C incubation chamber (Pecon). The hardware was controlled with the MetaMorph software (Molecular Devices). For classical time-lapse acquisitions, 300 to 600 sequential dual colour images of a single z plane 1 μm above the basal membrane were acquired every 250 ms with 491 nm and 561 nm lasers. For FRAP experiments, single colour acquisition with a 491 nm laser was performed on the basal membrane plane. 25 prebleach frames were acquired at a rate of 5 frames per seconds (fps). A 30 pixel circle region in the basal membrane was bleached by a double laser pulse without acquisition. Fluorescence recovery was followed by a stepwise acquisition of 2 s at a rate of 10 fps, then 1 min at 2 fps of and finally 9 min at 0.5 fps. The non-specific decay of the signal intensity in the bleached region due to the imaging process was corrected by the fluorescence decrease of the whole cell at each time point. The curves obtained were normalized so that the first post-bleach frame was set to zero and were then subject to curve fitting.

FRAP quantifications have been made using a standard modeling with chemical dominant interactions (Phair et al., 2004). First, raw intensity measurements were normalized using a double normalization in order to take into account the photobleaching and the background intensity. After the normalization, the intensities before and at the bleach time should reach respectively 1 and 0.

The recovery was estimated using the single exponential modeling

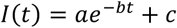

The parameters of equation have been estimated using a non-linear fitting within MATLAB (Mathworks) with the constraint *1*(*0*) = 0.

Once the parameters are estimated, the mobile fraction and the half-time recovery are given by: 
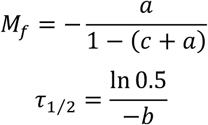
Each experiment was reproduced at least 3 times independently with a total sample number of at least 15 cells. Dynasore and hypotonic medium were added 2h and 10 min before the experiment, respectively.

### Stretching experiments

Briefly (a more detailed description of the device will be published elsewhere; Comelles and Riveline, in preparation), membranes and stencils were fabricated using polydimethylsiloxane (PDMS). PDMS (Sylgard 184 Silicon Elastomer, Dow Corning) polymer was prepared by mixing the base prepolymer with the cross-linking agent in a 10:1 ratio, vacuum atmosphere was used for 1h to remove gas bubbles from the mixture. The polymer was then poured over a plastic Petri dish and spin coated (Laurell WS-650-23B, Laurell Technologies) at 1000 rpm to achieve a 100 μm thick layer. Membranes were then cured for 12 hours at 65 °C. An analogous procedure was used to prepare the stencils: prepolymer was prepared at 10:1 ratio, degassed, poured over a Petri dish with a thickness of 1 mm and cured at 65 °C for 12 hours. 5 x 2.5 cm membranes and 2 x 2 cm stencils were cut off. Membranes were placed on a glass slide with parafilm. Stencils with vacuum grease (Dow Corning) on the bottom were placed on top of membranes. The setup was UV-sterilized under the cell culture hood and membranes were coated with poly-D-lysine mol wt 70,000-150,000 10 μg/ml in PBS for 1h at 37°C. Transfected cells were seeded on membrane and left to adhere for 24h. Imaging was performed on cells expressing EGFP-ITGB4(wt) and RFP-Arf6 or Cav1(wt or mutants) using an inverted Nikon Eclipse Ti equipped with HCX PL APO 63X NA 1.2 water objective (Leica), a Yokogawa CSU-X1 confocal head, an Evolve 512 EM-CCD camera (Photometrics), and a custom-made 37°C incubation chamber driven by the MetaMorph software (Molecular devices). Membranes were mounted on a custom-built cell-stretcher so that the membrane remained perfectly flat at the beginning of the experiment. An additional 10% uniaxial strain was applied on the cell stretcher by a computer controlled motor (Thorlabs). An image of each individual cell was taken at resting state, before applying the strain and a time-lapse was run at a rate of 1 image/min during 30 min under step stretch conditions.

### HD mechanoresponse analysis

Single plane images corresponding to the plasma membrane were analyzed using the *ImageJ* software. To quantify the area occupied by HDs, a ROI was manually drawn around each individual cell. For each cell, the first frame was used to determine the detection threshold. The same threshold was applied to the other frames.

HD growth and Cav1 disappearance between frame x and y was calculated as:

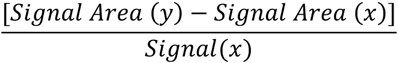
HD growth speed over 30 min was calculated as:

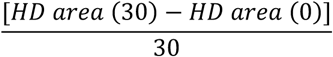
The final HD growth speed was computed as the median value of all individual cell values.

The orientation of HD growth was quantified using MATLAB (Mathworks). Appearing HD signal was segmented for each individual cell. Each cell was then centered on a polar coordinate system. Cells were subdivided in angular quadrants of π/16 radians. Angular histograms were computed as the proportion of overall intensity located in this angular portion. The final angular histogram is computed as an average of all cell's angular histograms.

### Osmotic shock

Cells were incubated for 10 min at 37°C with normal medium or with medium diluted in deionized water (1:9 for a 30 mOsm hypotonic shock). Cells were then either fixed, stained with antibodies specific for ITGA6 (HDs) and for Cav1 (caveolae) and their respective amount at the basal membrane was quantified from confocal images or used for live cell FRAP experiments as previously described.

### Transmission electron microscopy

Cells transfected in 6-well plates were fixed for 1h at room temperature with 2.5% glutaraldehyde and washed in 0.1M Na Cacodylate buffer. Cells were then post-fixed in 1% OsO4 for 30 min at 4°C, washed with 0.1M Na Cacodylate buffer, and finally with deionised water. Samples were secondary post-fixed in 4% water solution of uranyl acetate for 30 min at room temperature. The samples were stepwise dehydrated in ethanol (70% 20min, 3x95%, 3x100% each 15min) and infiltrated in diluted Epon (Ethanol/Epon=1/1) for 1h. Samples were left in absolute Epon (EmBed812) overnight. The following day, samples were placed in a fresh absolute Epon for 1h and left to polymerize at 60 °C for 24h - 48h. Once polymerized, samples were re-oriented and reembedded in empty epon blocks. Thin sections (100 nm) were sectioned using ultramicrotome (Leica Ultracut UCT), collected on formvar-coated slot grids and contrasted with 4% water solution of uranyl acetate for 10min and Reynolds lead citrate for 3min. Thin sections (100nm) were imaged with CM120 transmission electron microscope (Philips Bio Tween) operating at 120 kV. Images were recorded with Veleta 2k x 2k (Olympus-SIS) camera using iTEM software. Caveolae were manually counted on at least 20 pictures of a region of 3∼4 μm of plasma membrane in 20 different cells.

### Immuno-electron microscopy

Cells were grown to confluence in 75 cm^2^ flasks and chemically fixed with 2% paraformaldehyde + 0.2% glutaraldehyde in 0.1M Na Cacodylate buffer for 30min at room temperature, then fixed again with fresh fixative for 1h at room temperature. Cells were collected by 5 steps of centrifugation at 2500 rpm and resuspended in 1% glycin in PBS. The supernatant was replaced by liquid gelatin, cells were centrifugated for 3 min at 14000 rpm and incubated for 30 min on ice. Samples of size 1 mm^2^ were incubated overnight in 2.3M saccharose at 4°C under gentle agitation. Samples were cryo-fixed in liquid nitrogen and ultrathin sections (∼60nm) were cut with an EM FCS ultramicrotome (Leica). Samples were washed with PBS and then 1% glycin in PBS, saturated in 1% BSA in PBS for 15 min at room temperature and incubated with Cav1 antibodies (BD rabbit polyclonal) at dilution 1:50 in 1% BSA in PBS for 30 min. Samples were washed in 1% BSA in PBS, incubated with protein A coupled to 10 nm gold beads for 15 min at room temperature, washed with 1% BSA in PBS, and then with PBS. Samples were fixed in 1% glutaraldehyde in PBS for 5 min at room temperature and washed in PBS and then 1% glycin in PBS. Samples were saturated in 1% BSA in PBS for 15 min and incubated with ITGA6 antibodies (BD rat monoclonal) at dilution 1:50 in 1% BSA in PBS for 30 min, then washed in 1% BSA in PBS, incubated with anti-rat antibodies at dilution 1:200 in 1% BSA in PBS for 15 min at room temperature, washed with 1% BSA in PBS, incubated with protein A coupled to 15 nm gold beads for 15 min at room temperature and washed with 1% BSA in PBS and then PBS. Samples were fixed in 1% glutaraldehyde in PBS for 5 min at room temperature, washed in PBS and then deionised water and contrasted with 4% water solution of uranyl acetate and methyl cellulose on ice for 10min. Thin sections (60nm) were imaged with CM120 transmission electron microscope (Philips Bio Tween) operating at 120 kV. Images were recorded with Veleta 2k x 2k (Olympus-SIS) camera using iTEM software.

### Quantitative PCR with reverse transcription

Total RNAs of transfected cells were purified 72h after transfection with an RNeasy Midi kit (Qiagen). The cDNAs were produced using a SuperScript II reverse transcriptase and oligo(dT) primers (Invitrogen) following the manufacturer’s instructions. Quantitative PCR on cDNAs was carried out on a LightCycler 480 (Roche) with SYBR Green probes (Sigma). HPRT expression was used as an internal control for each reaction to normalize the cDNA input. The ratio of the expression level for each siRNA knockdown condition versus a scrambled siRNA control was then calculated. Primer sequences used: GIT1 forward CCATGGACGTGTATGACGAG, GIT1 reverse GCAAACTCTCGGGCA TTAAA, βPIX forward CCCAGGTCCTGATTCAGTGT, βPIX reverse CTGTAGGTGC TCCACCCATT, PAK1 forward AATCTGCCAGGGACATGAAG, PAK1 reverse CATTTTCCCCGACTTCTCAA, Rac1 forward AGCTTTTGCGGAGATTTTGA, Rac1 reverse CCCGTGACACTTTCATTCCT.

### Statistics

Student’s t-test was used for all the statistical analysis, except for GSD image colocalization, FRAP analysis, and TEM caveolae quantification where a Wilcoxon rank-sum test was used. All experiments were repeated at least 3 times independently.

---

**Figure S1.**
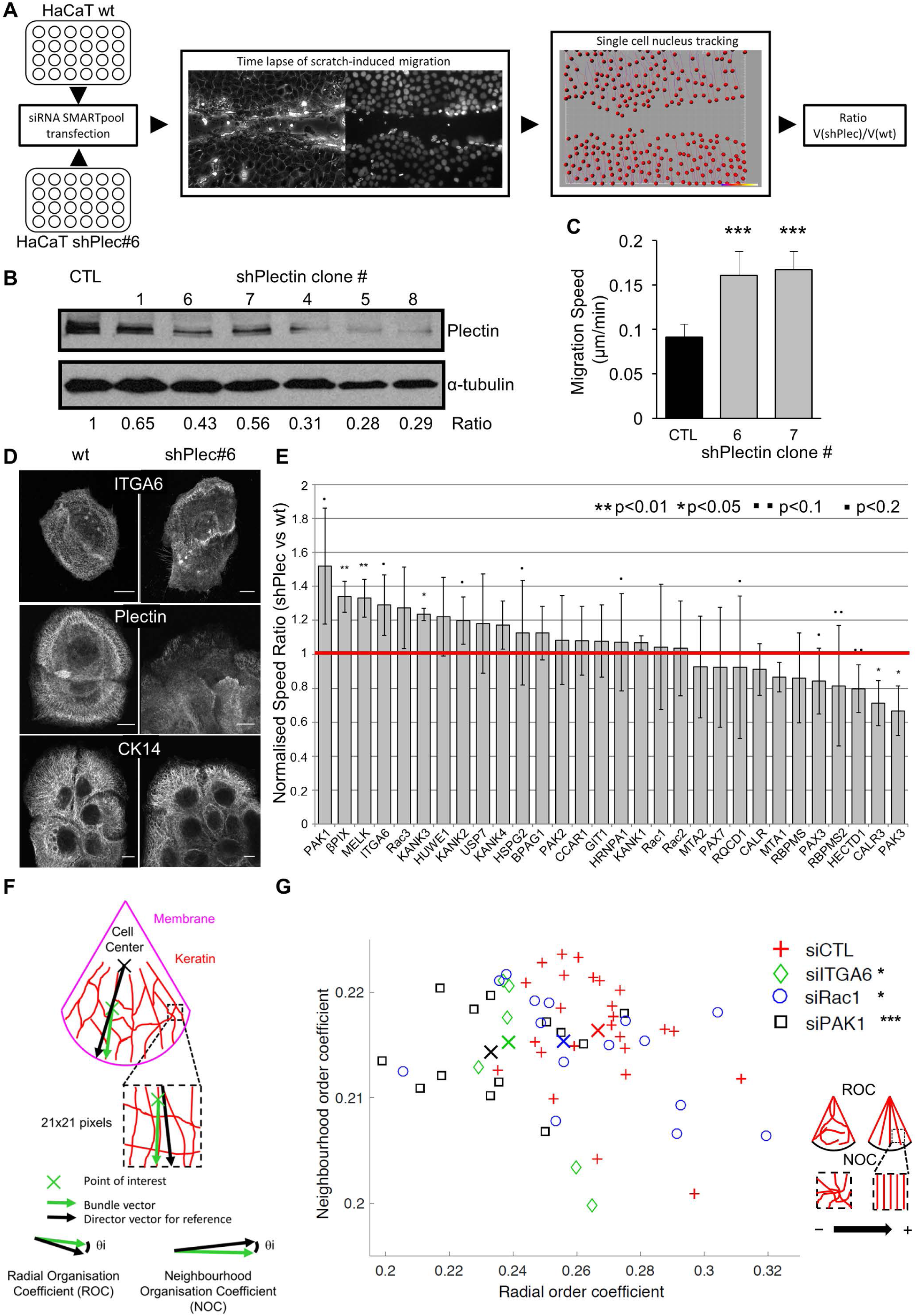
siRNA screen on HaCaT cells with reduced plectin levels. (A) Scheme describing the siRNA screening strategy used to identify regulators of HDs (see methods for details). (B) Protein extracts from independent HaCaT clones expressing an shRNA against PLEC mRNA were immunoblotted for plectin (top) and α-tubulin (bottom); protein expression was estimated by densitometric ratio. Uncropped films are shown in supplementary figure 9. (C) Wound healing migration of HaCaT-GFP-H1B wt and shPlectin clones was monitored for 24h and average speed of individual cells was measured over the whole migration. wt=3408, shPlec#6=3561, shPlec#7=3285 cells. (D) HaCaT-GFP-H1B wt and shPlectin#6 clones were stained for HD markers with ITGA6, plectin and CK14. Scale bar=10 μm. (E) The value [VshPlec/Vwt]_siGENE_/[VshPlec/Vwt]_SiCTL_ is shown for each targeted gene. Red line shows the threshold. (F) Scheme describing the method used to quantify the organization of the keratin network (see methods for details). (G) Quantification of keratin network organization in HaCaT cells transfected with control siRNAs or siRNAs against ITGA6, Rac1 or PAK1 (see methods for details).

**Figure S2.**
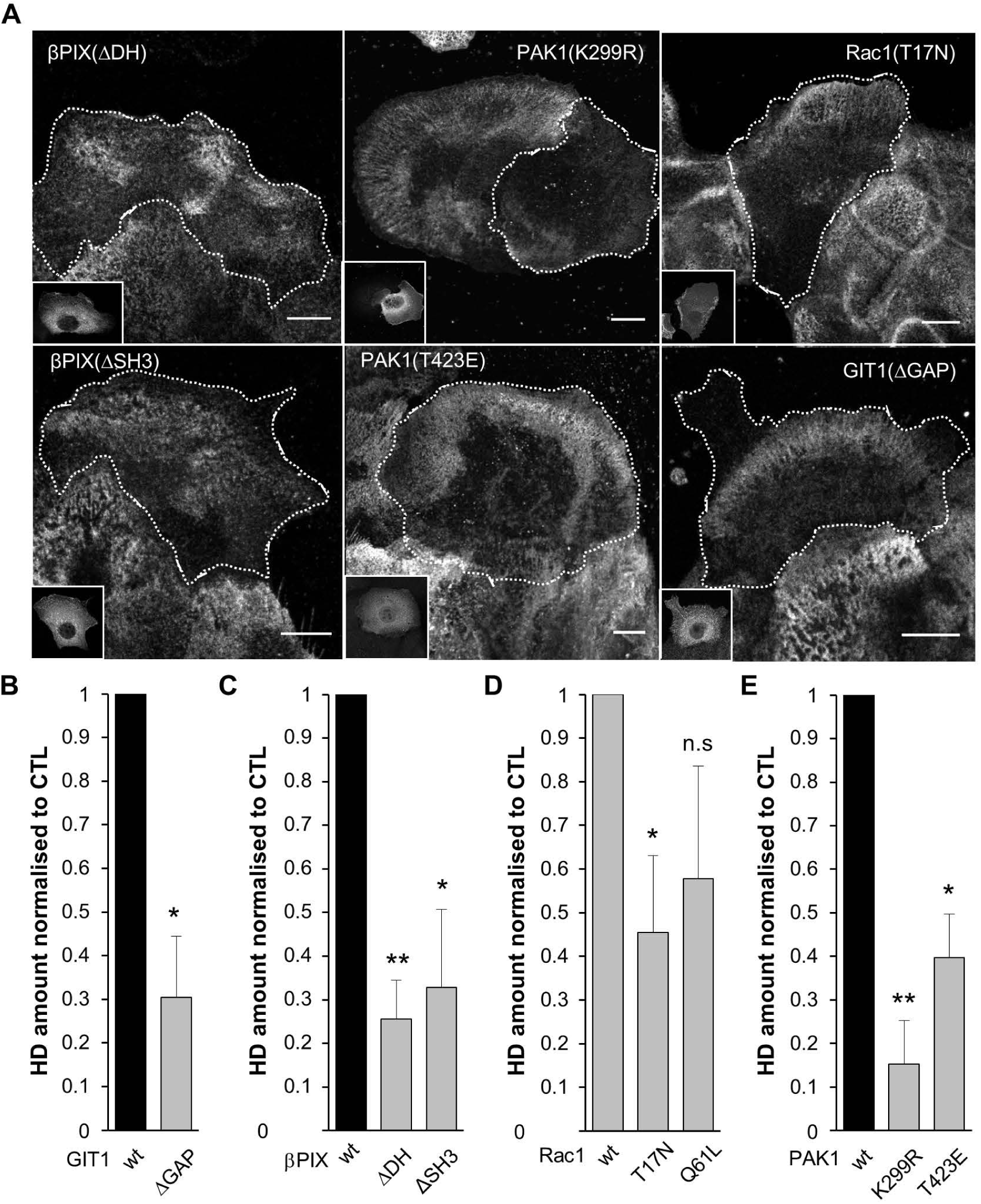
The protein-protein interaction domains and enzymatic activities of GIT1, βPIX, PAK1 and Rac1 are required for proper HD organisation. (A) Representative confocal images of the basal membrane of HaCaT cells transfected with mutant HA-βPIX, myc-PAK1, Flag-GIT1 and EGFP-Rac1 constructs. The following domains were tested: (i) the βPIX RacGEF (ΔDH) and SH3 (ΔSH3; interacting with PAK1) domains, (ii) the Rac1 GTPase and (iii) PAK1 kinase activities (dominant-negative Rac1(T17N) and kinase-dead PAK1(K299R), respectively). The inset on the lower left of each image shows the expression of each construct. Cells were stained for ITGA6 (HDs) and the indicated tag (except for EGFP). Scale bar=10 μm. (B-E) Quantification of the amount of HDs at the basal membrane of cells expressing (B) GIT1, (C) βPIX, (D) Rac1 or (E) PAK1 mutant forms (related to Fig. 1).

**Figure S3.**
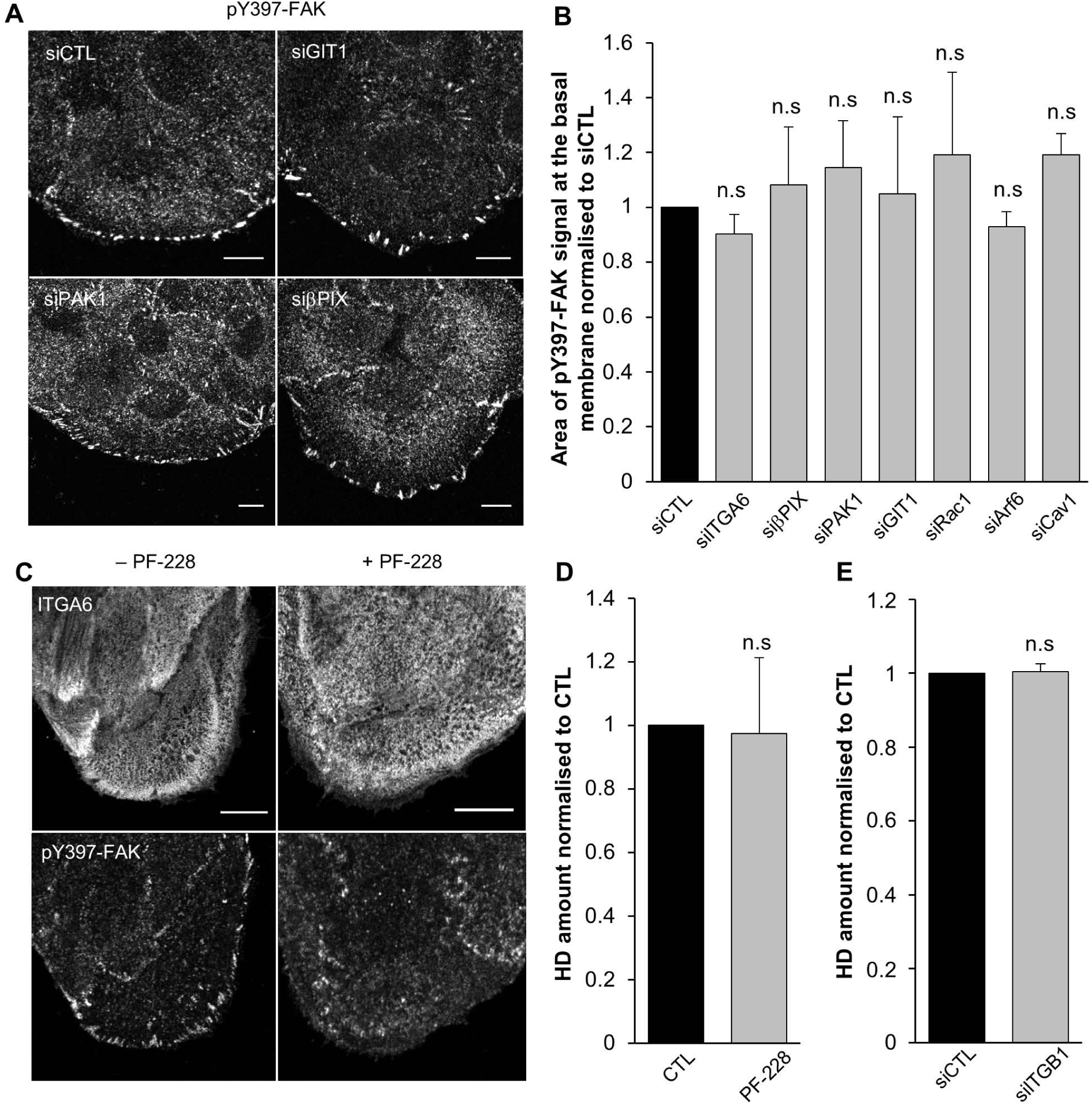
HD remodeling does not depend on FA signalling. Cells transfected with control or candidate genes siRNAs were stained for pY397-FAK. (A) Representative confocal images of the basal membrane of siRNA transfected cells stained for pY397-FAK immunostained for ITGA6 and pY397-FAK. Scale bar=10 μm. (B) The area covered by the signal relative to the overall cell area was measured and normalised to control siRNA. Note that the pY397-FAK signal at the basal membrane did not change suggesting that the various siRNA treatment did not indirectly affect HDs through a primary effect on FAs. (C) Representative confocal images of the basal membrane of cells treated with vehicle or the FAK inhibitor (PF-228) and immunostained for ITGA6 and pY397-FAK. Scale bar=10 μm. (D) Quantification of the amount of HDs on the basal membrane of cells treated with vehicle or the FAK inhibitor (PF-228). Note that affecting FAK activity with PF-228, did not induce any effect on HDs. (E) Quantification of the amount of HDs on the basal membrane of cells transfected with control siRNA (siCTL) or ITGB1 siRNAs (silTGB1).

**Figure S4.**
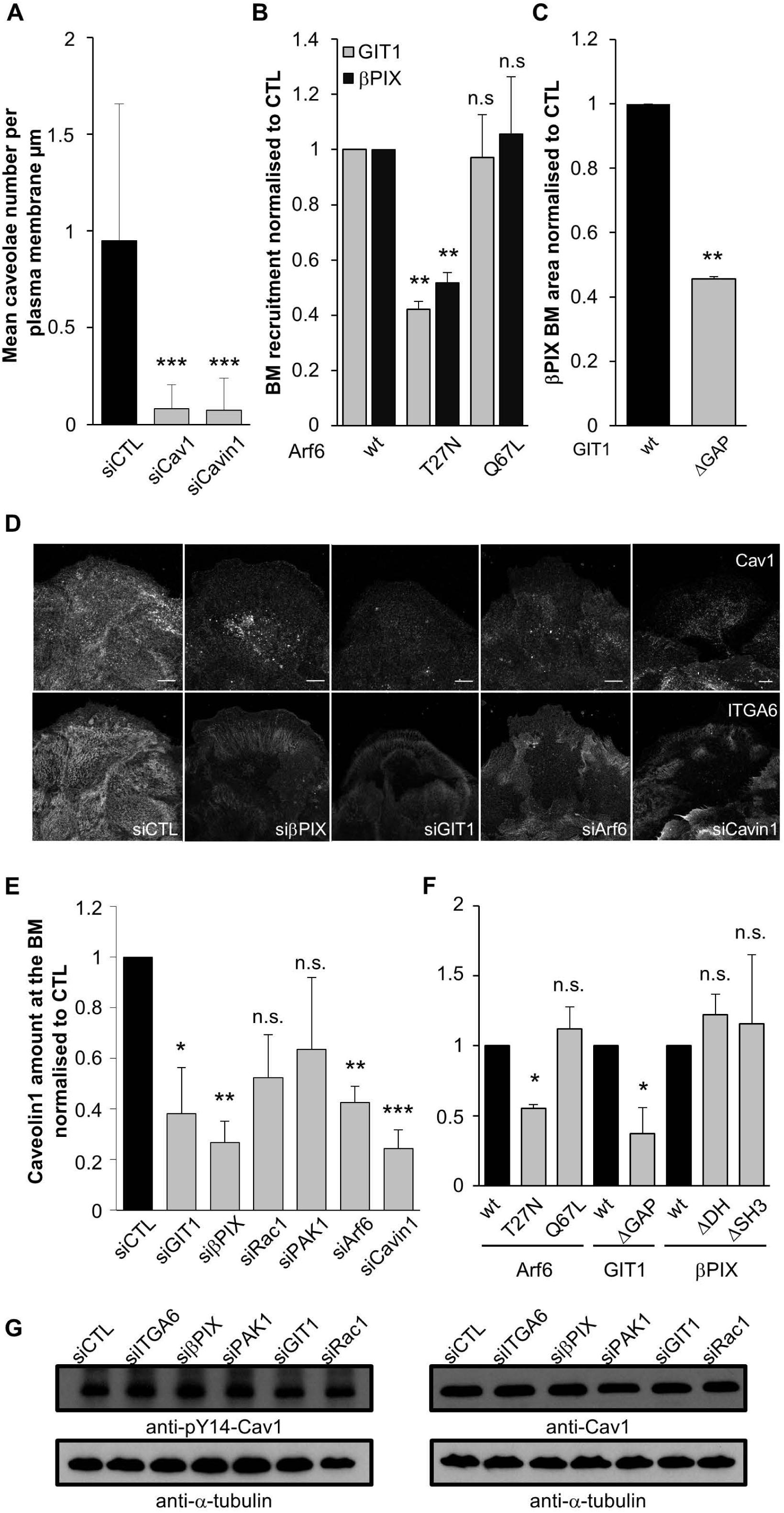
Arf6 and GIT1/βPIX are required for proper caveolin1 localisation at the basal plasma membrane. (A)Quantification of the mean caveolae number per μm of plasma membrane counted on transmission electron microscopy images of 100μm sections of cells transfected with control, anti-Cav1 or Cavin1 siRNAs (see methods for details). (B) Quantification of the area covered by βPIX (black) or GIT1 (grey) at the basal membrane of cells expressing EGFP-Arf6 mutant forms with Flag-GIT1wt (grey) or alone (black) and stained for Flag (grey) or βPIX (black). (C) Quantification of the area covered by βPIX at the basal membrane of cells expressing Flag-GIT mutant forms and immunostained for βPIX. (D) Confocal images of the basal membrane in cells transfected with control or indicated siRNAs and immunostained for ITGA6 and Cav1. Scale bar=10 μm. (E-F) Quantification of the area covered by Cav1 on the basal membrane of (E) siRNA-transfected cells, or (F) cells expressing Arf6, GIT1 or βPIX mutant forms. (G) Protein extracts of cells transfected with control, GIT1, βPIX, PAK1 or Rac1 siRNAs were immunoblotted with Cav1 or pY14-Cav1 antibodies and α-tubulin a as loading control.

**Figure S5.**
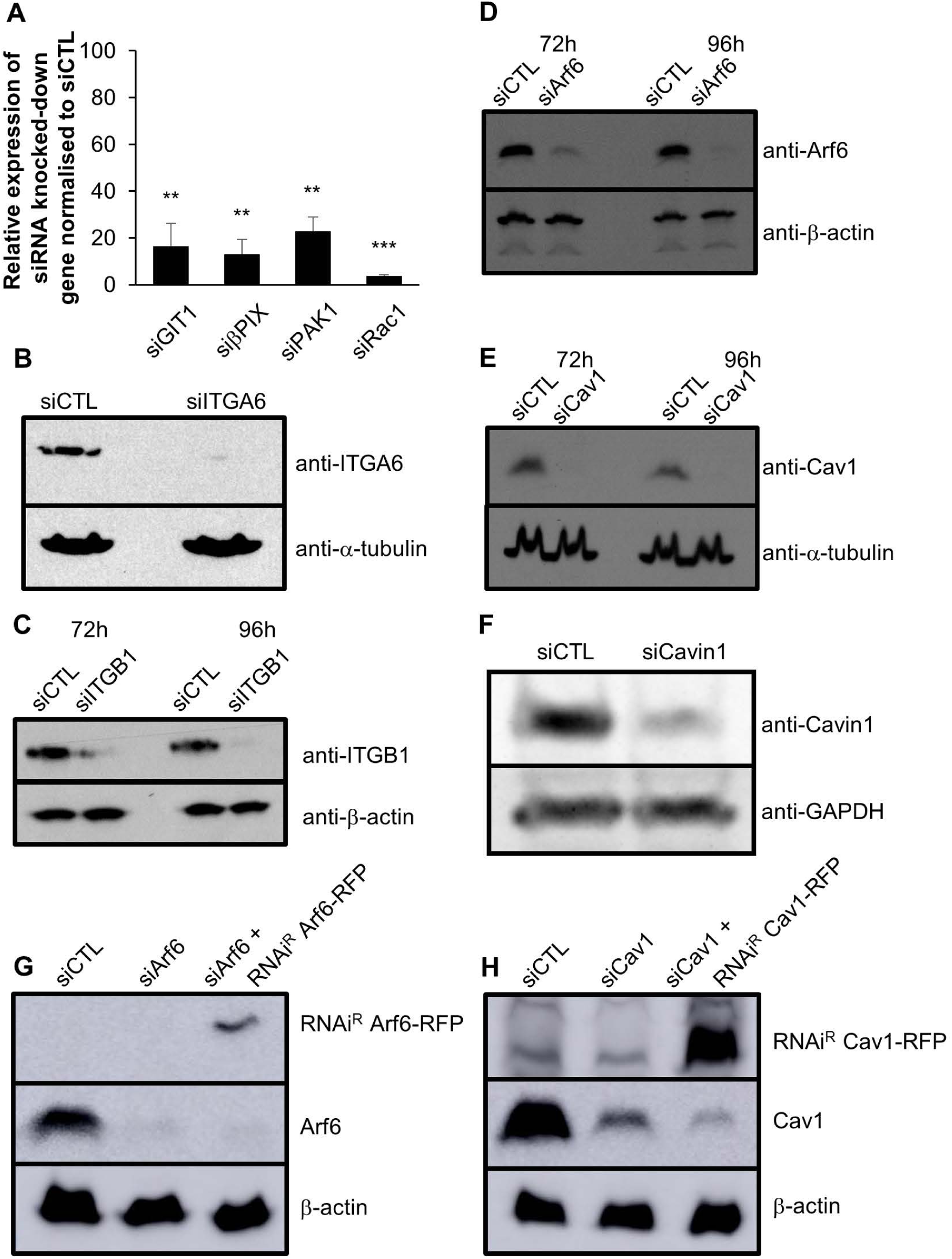
Quantification of siRNAs knockdown effect. (A) Total RNAs were extracted from cells transfected with control, GIT, βPIX, PAK1 or Rac1 siRNAs. RT-qPCR was performed pairwise on each control and knocked-down samples to assess the siRNAs efficiency. The ratio of gene expression (siRNA/siCTL) was calculated. (B-H) Protein extracts of cells transfected with control siRNAs or siRNAs against (B) ITGA6, (C) ITGB1, (D,G) Arf6, (E,H) Cav1 or (F) Cavinl were immunoblotted against (B) ITGA6, (C) ITGB1, (D,G) Arf6, (E,H) Cav1 or (F) Cavinl and (B,E) α-tubulin, (C-D,G-H) β-actin or (F) GAPDH as a loading control. For rescue experiments (G-H), cells were transfected with (G) Arf6 or (H) Cav1 siRNAs then 48h later nucleofected with the corresponding siRNA-resistant constructs (RNAi^R^) and protein extracts were prepared 24h later.

## Supplementary movie legends

**Video 1. ITGB4/Arf6 vesicles are transported between intracellular organelles.** Spinning disk confocal time lapse of a section +1 μm above the plasma membrane of a cell expressing EGFP-ITGB4 (green) and mCherry-Arf6(wt) (red). Images were acquired every 250 ms. The movie is played at 20 frame/s.

**Video 2. ITGB4/Cav1 vesicles are transported between intracellular organelles.** Spinning disk confocal time lapse of a section +1 μm above the plasma membrane of a cell expressing EGFP-ITGB4 (green) and RFP-Cav1(wt) (red). Images were acquired every 250 ms. The movie is played at 20 frame/s.

**Video 3. Arf6/Cav1 vesicles are transported between intracellular organelles.** Spinning disk confocal time lapse of a section +1 μm above the plasma membrane of a cell expressing EGFP-Arf6(wt) (green) and RFP-Cav1(wt) (red). Images were acquired every 250 ms. The movie is played at 20 frame/s.

**Video 4. HD grow under mechanical strain.** Spinning disk confocal time lapse of the basal plasma membrane plane of a cell expressing EGFP-ITGB4 (upper left/green) and RFP-Cav1 wt (upper right/red). Images were acquired every minute for 30 minutes. Bottom left shows HD appearing over time (see methods for details).

